# Antigenic cartography using variant-specific hamster sera reveals substantial antigenic variation among Omicron subvariants

**DOI:** 10.1101/2023.07.02.547076

**Authors:** Barbara Mühlemann, Jakob Trimpert, Felix Walper, Marie L. Schmidt, Simon Schroeder, Lara M. Jeworowski, Jörn Beheim-Schwarzbach, Tobias Bleicker, Daniela Niemeyer, Julia M. Adler, Ricardo Martin Vidal, Christine Langner, Daria Vladimirova, Derek J. Smith, Mathias Voß, Lea Paltzow, Christina Martínez Christophersen, Ruben Rose, Andi Krumbholz, Terry C. Jones, Victor M. Corman, Christian Drosten

## Abstract

SARS-CoV-2 has developed substantial antigenic variability. As the majority of the population now has pre-existing immunity due to infection or vaccination, the use of experimentally generated animal immune sera can be valuable for measuring antigenic differences between virus variants. Here, we immunized Syrian hamsters by two successive infections with one of eight SARS-CoV-2 variants. Their sera were titrated against 14 SARS-CoV-2 variants and the resulting titers visualized using antigenic cartography. The antigenic map shows a condensed cluster containing all pre-Omicron variants (D614G, Alpha, Delta, Beta, Mu, and an engineered B.1+E484K variant), and a considerably more distributed positioning among a selected panel of Omicron subvariants (BA.1, BA.2, BA.4/5, the BA.5 descendants BF.7 and BQ.1.18; the BA.2.75 descendant BN.1.3.1; and the BA.2-derived recombinant XBB.2). Some Omicron subvariants were as antigenically distinct from each other as the wildtype is from the Omicron BA.1 variant. The results highlight the potential of using variant-specifically infected hamster sera for the continued antigenic characterisation of SARS-CoV-2.

## Introduction

The SARS-CoV-2 pandemic has caused over 768 million infections and 6.9 million deaths as of June 2023 (*1*). The provision of vaccines greatly reduced infection fatality rates and disease burden (*2*). However, antigenic variability in SARS-CoV-2, facilitated by substitutions in the viral spike protein, resulted in the evasion of antibody reactivity acquired upon infection or vaccination (*3–6*).

In particular, SARS-CoV-2 Variant of Concern (VOC) Omicron and its subvariants arising from late 2021 show strong escape from serum neutralization in individuals infected with pre-Omicron variants such as the initial variant (Spike D614G), as well as variants Alpha, Beta, Gamma, and Delta (*7–13*). Omicron subvariants also escape antibodies generated by individuals vaccinated and boosted with wildtype-based vaccines (*7, 13*), which prompted the development and licensing of bivalent vaccine formulations containing additional antigens based on either the Omicron BA.1 or BA.4/BA.5 subvariants (*14, 15*). In late 2022, the circulating viruses were dominated by Omicron subvariants with convergent traits at several spike amino acid positions (346, 444-446, 450, 452, 460, 486, 490, 493), including the BA.5.-derived variants BF.7, BQ.1.18, several BA.2.75-derived variants, and recombinant lineages. Those variants show additional serum neutralization escape, raising concerns about the need to further update vaccines (*16–20*). Ongoing monitoring of circulating variants is inevitable for determining the need to update vaccine antigens.

In initial attempts to track SARS-CoV-2 antigenic evolution, human primary infection or vaccination sera were used to investigate antigenic differences between SARS-CoV-2 variants by cross-titration using different serum neutralization tests (*3–5, 7–10, 12, 21, 22*). The use of primary infection sera allows the measurement of the base antigenic distances between variants, i.e., serum neutralization titers that are not confounded by previous infections or vaccinations. Sera from individuals exposed to multiple variants may lead to an underestimation of antigenic distances (*23, 24*). With the exception of young children, large fractions of the population have now been infected or vaccinated (*25–27*). Therefore, obtaining sera from subjects with primary infections with new variants is becoming very difficult. For example, while it was still possible to identify unvaccinated individuals with first infections with Omicron BA.1 and BA.2 (*9, 10, 12*), doing so for Omicron BA.5 and future variants will be difficult. Likewise, identifying individuals with known infection histories (e.g., vaccinated with a breakthrough infection with a known variant) will also become more challenging as testing programs are discontinued.

An alternative is to use sera from experimental animals with serum antibody response characteristics similar to that of humans. Using experimental animal infections allows the generation of sera for any variant, with predefined infection histories, taken at standardized times after infection (*24*). Further, due to the higher titers resulting from experimental infections, animal sera would provide more ‘dynamic range’ as titers will not become undetectable as readily as in human convalescent sera tested against variants with large antigenic distance (*7, 28*). Syrian hamsters are a widely recognized animal model for COVID-19. Upon infection with SARS-CoV-2, these animals develop mild to moderate disease and show seroconversion (*28–31*). However, there is still a lack of experience with standardization of infection and assay conditions, and only one laboratory has developed and presented a complete approach that includes strain selection, immunization scheme, and serum neutralization evaluation (*28, 32*).

Here we provide an independent approach to Syrian hamster-based antigenic cartography. We use an efficient immunization scheme based on low- and subsequent high-dose intranasal inoculation and a refined neutralization assay criterion with titers determined on a continuous scale using a neutralization threshold of 90%. We characterize antigenic differences between six representative and one artificial pre-Omicron variants (D614G, Alpha, Beta, Delta, Mu, B.1+E484K) and eight Omicron variants (BA.1, BA.2, BA.4, BA.5, BF.7, BQ.1.18, BN.1.3.1, and XBB.2) using antigenic cartography (*33*). The Omicron variants BF.7 and BQ.1.18 are descendants of BA.5, having acquired one (R346T, BF.7) or three (R346T, K444T, N460K, BQ.1.18) additional substitutions in the spike protein. Both circulated with up to ∼10% or ∼35% prevalence globally, respectively. BN.1.3.1 is a descendant of BA.2.75, with an additional 10 substitutions in the spike protein compared to BA.2. XBB.2 arose from a recombination between two BA.2 lineages (BJ.1 and BM.1.1.1). The variant acquired 13 additional substitutions in the spike protein compared to BA.2, 11 of those identical to XBB.1.5, the globally most prevalent variant starting in February 2023. The results reveal considerable antigenic diversity among Omicron subvariants, which are often as antigenically distant from each other as BA.1 is from the pre-Omicron variants.

## Results

We infected 26 six week-old female Syrian hamsters with the D614G isolate B.1 (n=3), Alpha (n=2), Delta (n=3), Beta (n=3), an engineered B.1 virus containing the escape mutation E484K, termed B.1+E484K (n=3), Omicron BA.1 (n=3), Omicron BA.2 (two different isolates, BA.2 and BA.2-12, n=3 each), and Omicron BA.5 (n=3) (Table S1). To increase serum neutralization capacity and in an attempt to reduce disease burden, hamsters were infected twice, using a low virus inoculum for the initial infection, followed by a high-dose re-infection three weeks later. Sera were collected two weeks after the second infection.

Sera were titrated by plaque reduction neutralization test (PRNT) on Vero E6 cells against live virus isolates of D614G, Alpha, Delta, Beta, Mu, B.1+E484K, and Omicron subvariants BA.1, BA.2 (two isolates, BA.2 and BA.2-12), BA.4, BA.5, BF.7, BQ.1.18, XBB.2, and BN.1.3.1 (Table S1). The BF.7, BQ.1.18, BN.1.3.1, and XBB.2 variants were only titrated against the D614G, BA.1, BA.2, and BA.5 sera. The BA.5 sera were only titrated against the D614G and BA.1, BA.2, BA.5, BF.7, BQ.1.18, BN.1.3.1, and XBB.2 isolates. All other variants and sera were titrated all-against-all.

### Titer determination

We investigated different methods for inferring titers from plaque counts (Supplementary Text ‘Titer determination’). We tested the effects of using different sensitivity levels on the fold change and the antigenic map topology by inferring neutralization titers when 50, 75, 90, and 99% of all plaques were neutralized. Overall, the fold change estimated by the titers inferred with different neutralization criteria were comparable (fig. S2, S3). When inferring antigenic maps for each of the different neutralization criteria (fig. S4, S5), we found that in the maps from NT75 and NT90 titers, generally the Omicron BA.1 and BA.2 variants are positioned within the groups of BA.1 and BA.2 sera, respectively, whereas in the NT50 maps, these variants are positioned away from their homologous sera, suggesting that NT50 titers slightly overestimate the reactivity of those variants in relation to their homologous sera. We proceeded to use NT90 titers for the remaining analyses (Supplementary Text section ‘Titer determination’). In addition, we investigated whether discrete and continuous titers would allow us to draw similar conclusions with regards to measured fold change (fig. S6) and the inferred antigenic maps (fig. S7) and found fold changes and the antigenic maps to be similar between different methods (fig. S6, S7). In the following, we therefore refer to NT90 titers as determined using the neutcurve package (version 0.5.7) (*34*), constraining the lower end of the neutralization curve to zero (figs. S1-S8, Supplementary Text section ‘Titer determination’).

### Analysis of raw titers and fold change from homologous variant

All animals generated high antibody titers, with the highest titers for each serum ranging from 666 to >5120 (Fig. 1). Reactivity patterns among the hamsters infected with the same isolate were largely uniform (fig. S9). Titers between the two BA.2 isolates (fig. S10A), and between the BA.4 and BA.5 isolates (fig. S10B) were similar.

**Figure 1:**
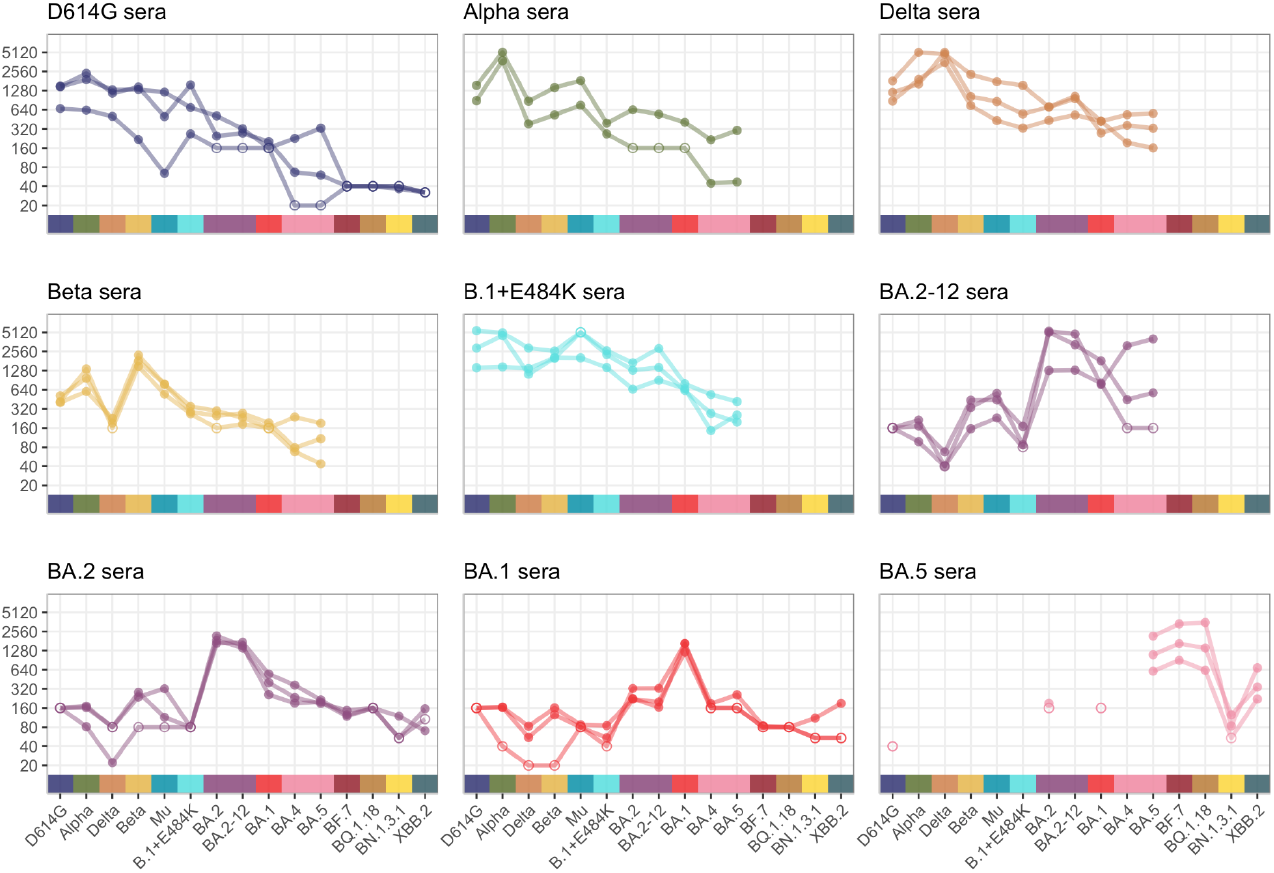
Neutralizing titers of 15 live virus isolates titrated against sera from hamsters twice infected with D614G, Alpha, Beta, Delta, Mu, B.1+E484K, Omicron BA.1, Omicron BA.2 (two isolates), and Omicron BA.5. Variants that were titrated are D614G, Alpha, Beta, Delta, Mu, B.1+E484K, Omicron BA.1, Omicron BA.2 (two isolates), Omicron BA.4, Omicron BA.5, Omicron BF.7, Omicron BQ.1.18, Omicron XBB.2, and Omicron BN.1.3.1. Detectable titers are shown as filled in circles, non-detectable titers are shown as empty circles. Since not all variant and serum pairs were tested in all dilutions, non-detectable titers range from <1:160 to <1:20.

In general, sera raised by infection with pre-Omicron variants have at least some reactivity to each non-homologous pre-Omicron variant, although the Beta sera have strongly reduced titers when titrated against the Delta variant (Fig. 1, 2). Against Omicron variants BA.1, BA.2, and BA.4/BA.5, titers of those sera tend to be lower, being highest against Omicron BA.2, and lowest against Omicron BA.4/BA.5. The D614G sera are not reactive or have low titers against Omicron variants BF.7, BQ.1.18, BN.1.3.1, and the XBB.2 recombinant. The sera raised by infection with Omicron variants show low or non-detectable titers against pre-Omicron variants. BA.1 and BA.2 sera retain some reactivity against BA.2 or BA.1, respectively, and have low or non-detectable titers against BF.7, BQ.1.18, BN.1.3.1, and XBB.2 Omicron variants. The BA.5 sera have high titers against the related BA.5, BF.7, and BQ.1.18 variants, and surprisingly, also show relatively high reactivity against the recombinant XBB.2, but not against BA.1, BA.2, or BN.1.3.1.

**Figure 2:**
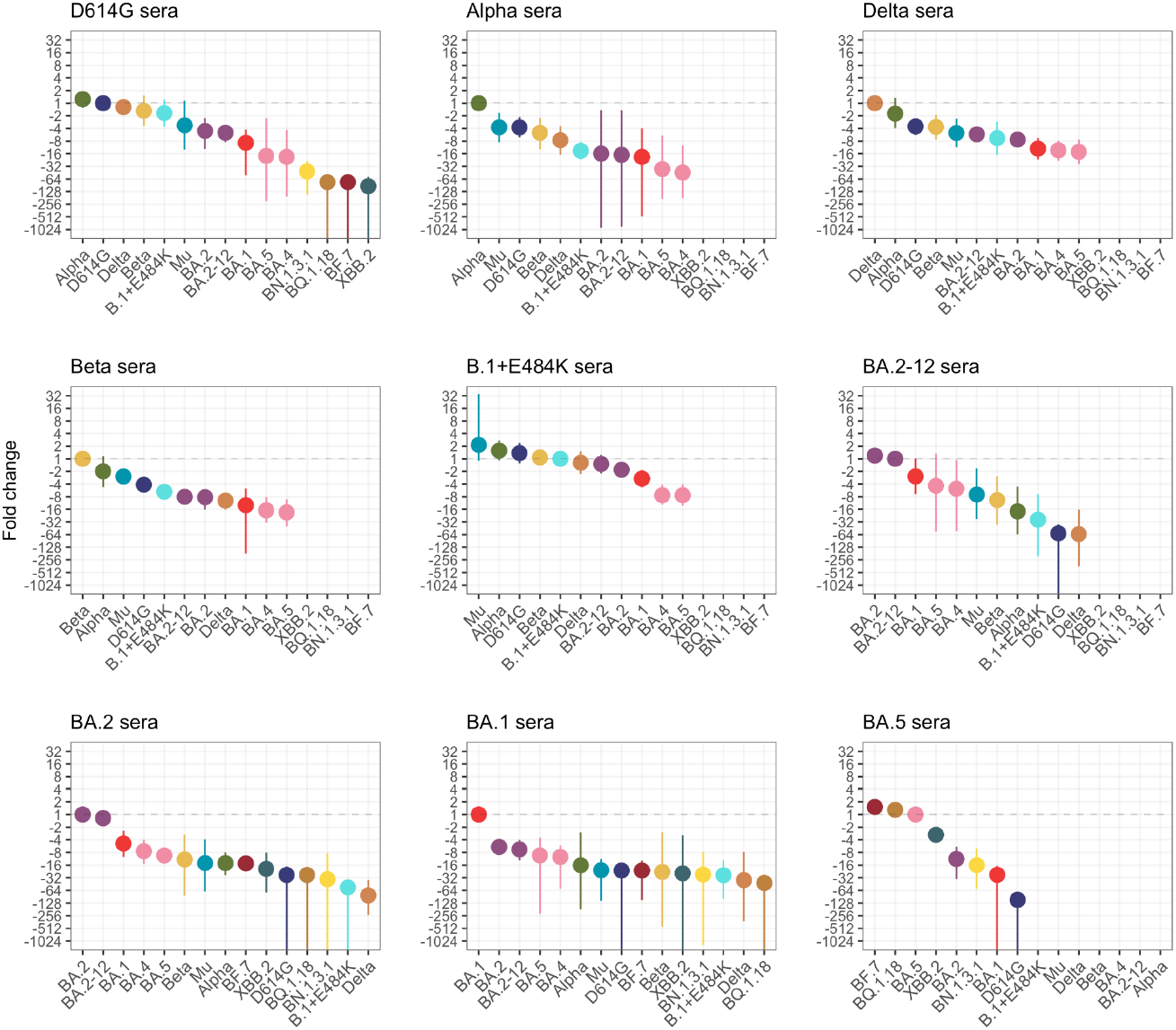
Fold change compared to the homologous variant. Circles show the point estimate for the mean fold change, while the intervals show the 95% highest posterior density interval of the fold change estimate. Variants are ordered by decreasing fold change within each serum group.

The raw titers and fold change show the considerable antigenic distance between pre-Omicron and Omicron variants. The differences in titers measured against different Omicron variants in Omicron sera also indicate a large antigenic diversity among Omicron variants.

### Antigenic cartography

We used antigenic cartography (*33*), implemented in the Racmacs package (*35*), to visualize the antigenic relationships between the different SARS-CoV-2 variants and sera. In the antigenic map, the distance between each variant and a serum corresponds to the fold-drop compared to the variant with the highest titer against that serum. The antigenic map (Fig. 3) provided a good fit to the data and was robust to missing titer data and noise (figs. S13-S25, Supplementary text ‘Assessing antigenic map model fit’). Antigenic maps can in principle be constructed in any number of dimensions. While a higher number of dimensions can lead to a better fit to the data, it can, like any statistical method, result in overfitting. Consequently, if the fit in higher dimensions are similar, then the lower dimensional representation is preferred, and is also easiest to visualize and interpret. Using cross-validation, we found that two dimensions provided an adequate fit to the data (fig. S14), although the BN.1.3.1 variant was placed closer to the BA.1 and D614G variants in the three-compared to the two-dimensional antigenic map (fig. S14).

**Figure 3:**
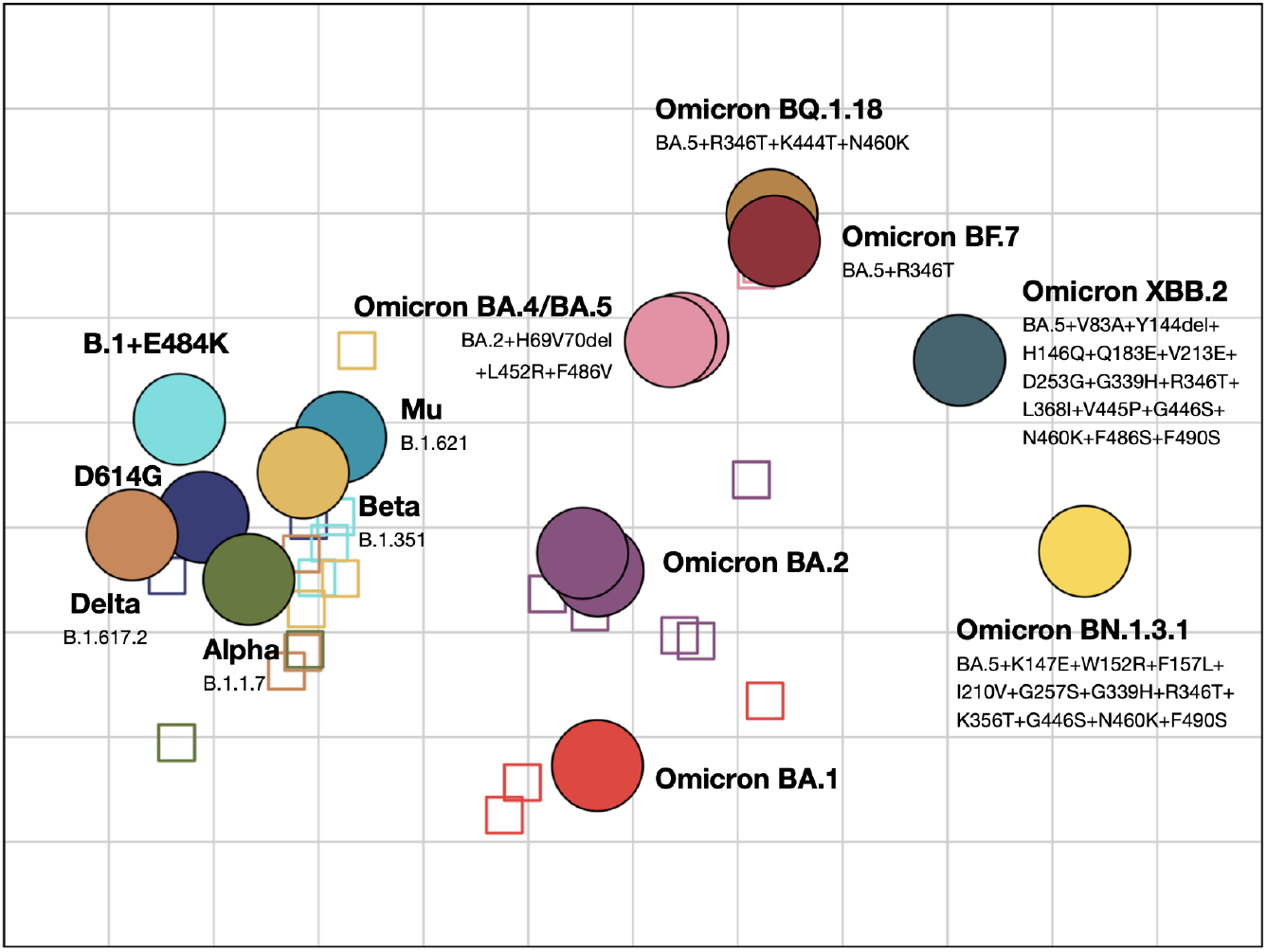
Antigenic map showing antigenic relationships between SARS-CoV-2 variants and sera. Distances between each variant and serum correspond to the fold change to the maximum titer for each serum. Viruses are shown as circles, sera as squares, with sera coloured by the color of their homologous variant (blue: D614G, green: Alpha, dark-yellow: Beta, orange: Delta, green-blue: Mu, cyan: B.1+E484K, red: BA.1, orchid: BA.2 (2x, on top of each other), pink: BA.4 and BA.5, ochre: BQ.1.18, maroon: BF.7, sea-green: XBB.2, yellow: BN.1.3.1). The side length of each grid square corresponds to a two-fold serum dilution in the neutralization assay. The rotation of the map is arbitrary, and is here oriented to correspond to previously published maps (*7, 9, 28*).

The antigenic map (Fig. 3) of the titers presented in Fig. 1 shows a cluster consisting of pre-Omicron variants (D614G, Alpha, Delta, Beta, Mu, B.1+E484K), and a looser clustering of Omicron subvariants, consistent with the titer and fold-drop pattern observed in Figs. 1 and 2. Within the Omicron subvariants, BA.4/BA.5, BF.7 and BQ.1.18 group relatively closely together, in agreement with the descent of BF.7 and BQ.1.18 from BA.5. The map shows larger antigenic distances between the other Omicron variants than between the pre-Omicron variants. Interestingly, the larger antigenic distances in the antigenic map are mainly caused by the inclusion of titrations against the BA.1, BA.2 and BA.5 sera, as excluding those serum groups moves the Omicron variants closer together (fig. S21). This indicates there are differences among the Omicron variants that are detected by BA.1, BA.2 and BA.5 sera, and not by the pre-Omicron sera we have used in this study. In agreement with the raw titer data, the antigenic map shows considerable antigenic diversity among the Omicron variants, with many Omicron variants (e.g. BA.1 vs BA.4/5, BA.4/5 vs BN.1.3.1) being as antigenically distant from each other as they are from the pre-Omicron variants.

To limit the future use of animals needed, we investigated the minimum number and type of sera required for appropriate antigenic map triangulation by subsampling the sera included in the map. For this analysis, we considered only those virus isolates and sera that had been included in all titrations, so that each variant and serum is positioned by the same number of titrations. Therefore, we excluded titrations of the Omicron BA.5 sera and BF.7, BQ.1.18, XBB.2, and BN.1.3.1 variants that have not been titrated against all variants or sera in the map, respectively. We constructed antigenic maps from randomly subsampled combinations of sera and serum groups (figs. S19, S26, S27). Both BA.1 and BA.2 serum groups are required for adequate positioning of the Omicron variants (fig. S19, S27A,B,C). To determine the number of pre-Omicron serum groups required for positioning the antigens and sera on the antigenic map, we subsampled from the pre-Omicron serum groups and always including the BA.1 and BA.2 sera. This resulted in antigenic maps that were largely similar to the map made from all sera (Fig. 3), even when only one pre-Omicron serum group was included alongside the BA.1 and BA.2 serum groups (fig. S22, S26, S27C). We further subsampled maps made from three to seven serum groups to contain only one or two sera per serum group (fig. S27D,E), and found the antigen positions of those maps to be more different to the full map compared to the maps made from all sera within a serum group (fig. S27C). This suggests that further reducing the number of sera within a serum group negatively affects the topology of the antigenic maps. However, the number of different pre-Omicron virus variant serum groups can be decreased.

## Discussion

Here we present an antigenic map of SARS-CoV-2 variants using sera from hamsters infected twice with viruses selected from a set of eight SARS-CoV-2 variants. The antigenic map shows a cluster consisting of the pre-Omicron variants and distinct positions of the Omicron variants. The map is qualitatively similar to maps made with human sera but reveals substantial antigenic diversity, in particular among Omicron subvariants (*7–10, 12*).

When others have compared human serum titers of triply vaccinated individuals with those of people with the same vaccination history and additional BA.1 or BA.2 breakthrough infections, the fold-drop between BA.1 and BA.4/BA.5 is usually larger in the BA.1-than the BA.2 breakthrough infectees and the thrice vaccinated (*36–41*). These results are consistent with our findings that show marked antigenic differences between Omicron variants that are not necessarily obvious in pre-Omicron sera. Interestingly, breakthrough infections with BA.5 after three vaccinations results in a broader immune response to BA.1, BA.2, and BA.4/BA.5 (*38, 41*), contrasting with sera from BA.5-infected hamsters, which reveal a clear differentiation of Omicron BA.1, BA.2, and BA.4/5 variants (*32, 41, 42*).

For a robust placement of variants in an antigenic map, it is advantageous if the sera are antigenically diverse. This is highlighted by the finding that the antigenic diversity between the Omicron subvariants is much less visible on maps made from pre-Omicron sera only (fig. S21A-C). Generally, it was possible to reduce the number of serum groups in an antigenic map consisting of the pre-Omicron and early Omicron (BA.1, BA.2, and BA.4/5) variants to as few as three, provided that the three variants against which the sera were raised were antigenically diverse. However, a reduction in serum group size to less than three sera led to increased distortion of the antigenic map. To reduce the number of experimental animals, an informed selection of immunization strains rather than reduction of animal group size is advisable.

The antigenic map shown here is similar to the map published by Mykytyn et al., 2023, which also uses hamster sera (*28, 32*). Exceptions are the placement of the BA.5 variant closer to BA.2, possibly caused by the absence of sera raised against the BA.2 variant in that study, and the different placements of the XBB.1 and XBB.2 variants in the two maps (these variants only differ at positions 252 and 253 of the spike protein). The hamster sera in Mykytyn et al. were raised by infecting hamsters once, with a dose of either 1.0×10^5^ or 5.0×10^4^ PFU, while the hamsters used here were infected twice with a priming dose of 1.0×10^3^ PFU followed by a booster dose of 1.0×10^6^ PFU. A prime-boost immunization regimen may alter mature antibody specificity and affinity, resulting in antigenic map differences.

When compared to first-infection human sera, the maps based on hamster sera presented here and in (*32*), agree in their tighter clustering of the pre-Omicron variants and larger differences between the Omicron subvariants (*7–10, 12*). The human sera maps show larger relative differences among the pre-Omicron variants, and a tighter placement of the Omicron variants. The tighter clustering of the pre-Omicron variants in the hamster sera data may be due to a more standardized immunization regimen in hamsters. Furthermore, the human sera antigenic maps have been generated predominantly employing lentivirus pseudotype or Focus Reduction Neutralization assays, while the hamster maps used PRNTs and authentic SARS-CoV-2 variants. As the antigenic distance between the Omicron variants is mainly governed by Omicron sera, the smaller distances observed among the Omicron variants in human sera maps may be caused by the smaller number of Omicron than pre-Omicron sera in those maps, as well as generally lower titers generated by infection with those variants in infection-naive humans.

Our study has the following potential limitations. First, the three hamsters infected per variant may not depict the full variability of individual responses to the different variants. We cannot assess here to what extent variability of individual responses in hamsters reflects that of humans. Second, while the antigenic map presented here is broadly similar to other maps, there are some differences. For example, the higher than homologous titers of BF.7 and BQ.1.18, as well as the higher titers of XBB.2 compared to BN.1.3.1 in BA.5 sera disagree with data published by Mykytyn *et al* (*32*). Some of the variability may be caused by the use of different live virus isolates in different laboratories, in particular substitutions affecting furin cleavage (*43*). Also, we performed titrations on VeroE6 cells that do not express TMPRSS2, which may lead to slightly different neutralization patterns in variants favoring TMPRSS2 dependent entry (even though neutralization mainly depends on receptor engagement) (*44*). Variations between datasets highlight the importance of performing antigenic surveillance in multiple laboratories using different assays.

With the majority of the population possessing a variable degree of immunity against SARS-CoV-2, there is a clear and pressing need to develop an animal first-infection sera model system to measure SARS-CoV-2 antigenic variation. The data presented here underscore the suitability and utility of using mono-specific hamster sera. Continually extended antigenic maps will be invaluable for maintaining an understanding of SARS-CoV-2 antigenic evolution and will aid vaccine strain selection decisions.

## Materials and methods

### Cell cultures and viruses

Vero E6 (ATCC CRL-1586) and Calu-3 (HTB-55) cells were maintained at 37°C, 5% CO2 in culture medium Dulbecco’s Modified Eagle’s Medium (DMEM, ThermoFisher Scientific) supplemented with 10% fetal bovine serum (FBS, ThermoFisher Scientific), 1% non-essential amino acids (ThermoFisher Scientific) and 1% sodium pyruvate 100 mM (NaP, ThermoFisher Scientific). All cell lines tested negative for Simian virus 5 (*Orthorubulavirus mammalis*) and mycoplasma contaminations. Low passage virus stock solutions were generated in Vero E6 or Calu-3 cells and sequenced to investigate the presence of substitutions introduced during isolation and passaging. Results are given in Table S1. The B.1+E484K virus was generated using the transformation-associated recombination (TAR) cloning method to introduce an E484K exchange into an infectious SARS-CoV-2 cDNA clone (*45, 46*).

### Animal husbandry

Six-to 10-week-old female Syrian hamsters (*Mesocricetus auratus*; breed RjHan:AURA) were purchased from Janvier Labs and kept in groups of 1-3 animals in individually ventilated cages (GR900; Tecniplast). Animals were provided with bountiful enrichment (Carfil) and had unrestricted access to food and water at all times. Cage temperatures ranged from 22 to 24°C and relative humidity ranged from 40 to 55%. All animal work was performed in a certified BSL-3 facility. Hamsters arrived at the facility 7 days prior to the start of the experiments.

### Generation of hamster sera

For each SARS-CoV-2 variant (D614G, Beta, Delta, B.1+E484K, Omicron BA.1, Omicron BA.2 (two isolates), and Omicron BA.5, Table S1), three hamsters were intranasally infected to generate defined immune sera directed against an individual virus variant. Two individuals were infected with the Alpha variant. To minimize infection-induced distress and in the same time maximize serum antibodies, hamsters were infected using a two-stage protocol. Each animal was initially infected with 1000 plaque forming units (PFU) of the respective variant and allowed to recover from infection for 21 days. On day 21 after initial infection, animals were re-infected with 1×10^6^ PFU of the same variant. All infections were carried out intranasally under general anesthesia and in a total volume of 60 μl as previously described (*47*). 14 days after re-infection, animals were euthanized and blood was collected for serum harvest. During the experiment, all hamsters were monitored twice daily for clinical signs of disease in compliance with a state-authority approved animal use protocol (Landesamt für Gesundheit und Soziales in Berlin, Germany, approval number 0086/20).

### Plaque reduction neutralization tests

The neutralizing activity of each serum against the different virus strains was determined by plaque reduction neutralization test (PRNT), largely as described before (*48*) (Wölfel et al., 2020). In short, Vero E6 cells (1.6×10^5^ cells/well) were seeded in 24-well plates and incubated for ∼24 hours. Hamster sera were serially diluted in OptiPro medium and mixed with medium containing 100 PFU of the respective virus, incubated at 37°C for one hour and then added to the Vero E6 cells in duplicate. After a further hour at 37°C, supernatants were discarded, the cells washed once with PBS and supplemented with 1.2% Avicel solution in DMEM. After three days at 37°C, the supernatants were removed, the plates were fixed using a 6% formaldehyde/PBS solution and stained with crystal violet. All serum dilutions were tested in duplicates. Plaques were counted for each well. The virus strains used were, Munich isolate 984, D614G (GISAID ID: EPI_ISL_406862), Alpha (EPI_ISL_754174); Beta (EPI_ISL_862149); Delta (EPI_ISL_3710429); Omicron BA.1 (EPI_ISL_7019047); Gamma (kindly provided by Dr. Chantal Reusken via EVAg); Omicron BA.2 CSpecVir26729_2 (EPI_ISL_9553926); Omicron BA.2 CSpecVir26729_12 (EPI_ISL_9553935); Omicron BA.4 (EPI_ISL_16221624); Omicron BA.5 (EPI_ISL_16221625), Omicron BF.7 (EPI_ISL_17293602), Omicron BQ.1.18 (EPI_ISL_17293256), Omicron XBB.2 (EPI_ISL_17548230), Omicron BN.1.3.1 (EPI_ISL_17549547).

### Analysis of neutralization titers

We investigated different methods for inferring titers from the raw plaque count data. This included assessing the effects of using different plaque reduction criteria / sensitivity levels (NT50, NT75, NT90, or NT99 titers) (figs. S1-S5), as well as assessing the effect of estimating titers either by rounding to the closest dilution (‘discrete titers’), or by fitting a sigmoid neutralization curve (‘continuous titers’) (figs. S6, S7). Discrete titers were determined as the dilution closest to where X% of all plaques are neutralized, where X corresponds to the sensitivity level (e.g. 50 for NT50 titers). Continuous titers were inferred using the `HillCurve` function in the neutcurve package, version 0.5.7 (*34*) with and without constraining the lower and / or upper ends of the neutralization curve to zero or one, respectively (fig. S1).

For the analysis of reactivity patterns of different serum groups and for antigenic cartography, NT90 titers were determined from the raw plaque counts using the neutcurve package (version 0.5.7) (*34*) written in Python, constraining the lower end of the neutralization curve to zero (Supplementary Text section ‘Titer determination’). We used titers determined with the lower and upper end of the neutralization curve constrained at zero and one, respectively for the 28 pairs of sera and variants listed in Table S2. Geometric mean titers (GMT) and fold changes were estimated using the `gmt` and `log2diff` functions, respectively, in the titertools package (*49*) in R (version 4.2.0) (*50*), as described in (*7*). We used the following parameters: ci_method=‘HDI’ (highest density interval), ci_level=0.95, dilution_stepsize=0, the prior for the mean was a normal distribution with mean of 0 and standard deviation of 100, the prior for the standard deviation was an inverse gamma distribution with a shape parameter of 2 and a scale parameter of 0.75.

### Antigenic cartography

Briefly, the target distance in an antigenic map optimization between each variant and serum pair is inferred by calculating the difference between the log_2_ highest overall titer of the serum and the log_2_ titer of a given variant. Therefore, the variant with the highest titer against a particular serum will have a target distance of 0. The positions of the variants and sera in the antigenic map are adjusted in an optimization process that aims to minimize the sum of the squared differences between the target distances and the corresponding distances in the antigenic map (*33*). Antigenic maps were inferred using Racmacs version 1.1.39 (*35*). All antigenic maps were constructed with a dilution_stepsize of 0 for maps made from continuous titers or 1 for maps made from discrete titers. The maps were optimized in two dimensions for 500 optimizations, and the minimum column basis parameter set to “none”.

We performed several analyses to assess the dimensionality of the antigenic map (fig. S14), how well the distances in the antigenic map represent the target distances (figs. S15, S16), the robustness of the map to titer noise (fig. S17) and missing titers (figs. S18-23), and the predictive power of the antigenic map (figs. S24-S26), described in Supplementary Text section ‘Assessing antigenic map model fit’.

## Acknowledgements and funding sources

This work was funded by the German Federal Ministry of Education and Research through project DZIF (8040701710 and 8064701703) and VARIpath (01KI2021), as well as the German Federal Ministry of Health through project SeroVarCoV.

CD received additional funding from ECDC project Aurorae (NP/21/2021/DPR/25121) and EU Hera project Durable (101102733).

DJS and TCJ are part funded through the NIH NIAID Centers of Excellence for Influenza Research and Response (CEIRR) contract 75N93021C00014 as part of the SAVE program. VMC is a participant in the BIH-Charité Clinician Scientist Program funded by Charité - Universitätsmedizin Berlin and the Berlin Institute of Health.

VMC has his name on patents regarding SARS-CoV-2 serological testing and monoclonal antibodies.

