## Supplementary Information for "Antigenic cartography using variant-specific hamster sera reveals substantial antigenic variation among Omicron subvariants"

#### Supplementary Text

##### Titer determination

Effects of using discrete and continuous titers

Visual inspection of the neutralization curves in fig. S1 showed that continuous titers inferred when fixing the neutralization curve at zero provide the most parsimonious fit to the data. Given the overall similarity of the fold change and antigenic maps estimated by the different methods, we therefore proceeded to use continuous titers inferred when fixing the neutralization curve at zero. However, visual inspection of the neutralization curves in fig. S1 also indicated 28 titers (2.1 vs Alpha, 2.2 vs BA.2_2, 3.2 vs BA.2_12, 3.3 vs Alpha, 3.3 vs Beta, 4.1 vs Alpha, 4.2 vs Delta, 4.3 vs Alpha, 4.3 vs Delta, 5.1 vs Alpha, 5.1 vs BA.2_12, 5.2 vs Alpha, 5.2 vs BA.2_2, 5.3 vs BA.2_2, 6.1 vs BA.2_2, 6.2 vs BA.2_2, 7.1 vs D614G, 7.1 vs Alpha, 7.1 vs Delta, 7.1 vs E484K, 7.1 vs BA.2_12, 7.1 vs Mu, 7.2 vs Mu, 7.3 vs Alpha, 7.3 vs Mu, 8.1 vs BA.2_12, 9.1 vs BF.7, 9.3 vs BQ.1.18) where continuous titers inferred when fixing the lower and upper ends of the neutralization curve at zero and one, respectively provided a better fit to the data. These titers were therefore adapted as shown in Table S2. Antigenic maps with and without these adapted titers only show minimal differences (fig. S8, RMSD = 0.12).

##### Assessing antigenic map model fit

We performed several analyses to assess the dimensionality of the antigenic map (fig. S14), how well the distances in the antigenic map represent the target distances (figs. S15, S16), the robustness of the map to titer noise (fig. S17) and missing titers (figs. S18-23), and the predictive power of the antigenic map (figs. S24-S26).

In order to assess the dimensionality of the antigenic map, we investigated how well the antigenic distances in maps optimized in different dimensions fit the measured titers. To this end, we performed 1000 repeats where 10% of the titers were excluded at random, the map re-optimised using 500 optimisations in 1-5 dimensions, and calculated the root-mean-squared error between the measured titers and the predicted titers in the antigenic map. The map optimized in four dimensions fit the data best (RMSE of detectable titers = 1.056), but only performed marginally better than maps optimized in two (RMSE of detectable titers = 1.11), three (RMSE of detectable titers = 1.069), and five (RMSE of detectable titers = 1.059) dimensions (fig. S14C). The overall arrangement of the variants in a map optimized in three dimensions closely resembles the arrangement of variants optimized in two dimensions (fig. S14A, B), with the exception of the Omicron BN.1.3.1 variant, which in 3D takes up a position closer to the BA.1 and D614G variants. For ease of representation, we show the map in two dimensions.

We next assessed how well the distances fitted in the antigenic map represent the measured distances inferred from the titers (figs. S15, S16). Overall, there was a good fit between the fitted and the measured titers. The difference between the fitted and measured titers was

0.04 on the $\log_2$ scale, with a standard deviation of 0.73. When assuming a mean of 0, the standard deviation is 0.73. The standard deviation was higher than the standard deviation observed between repeated titrations (sd = 0.46) (fig. S13). However, because the repeats were not independent, the repeat variation may be underestimated, and the difference between fitted and measured titers is comparable to that observed in other studies investigating SARS-CoV-2 antigenic variation (*2*, *3*). We also found no evidence of increased differences between fitted and measured titers for specific serum groups and variant pairs (fig. S16).

We next investigated the robustness of the map to measurement error, variant reactivity biases and missing data. To assess robustness to measurement error and variant reactivity biases we performed a bootstrap analysis where normally distributed noise was added to the titer and / or variant reactivity (fig. S17). We performed 1000 bootstrap repeats of adding random noise and summarized the positional variation of the variants and sera, such that the areas shown in fig. S17 represent 68% (one standard deviation) of the positional variation of each variant or serum. Random noise added to the titers had a standard deviation of 0.35, as estimated from the standard deviation of differences between repeat titrations (fig. S13). Random noise added to variant reactivity had a standard deviation of 0.4, following Wilks et al., 2022 (*2*). In general, positions were largely robust to the addition of random noise, in particular for the Omicron BA.1, BA.2, BA.4/5, BF.7, and BQ.1.18 variants, and larger positional uncertainty within the pre-Omicron variants and for Omicron XBB.2 and BN.1.3.1. Next, we assessed the robustness of the antigenic map to the removal of each variant (fig. S18) and serum group (fig. S19) in turn. Variant positions were largely robust to the removal of individual variants, with increased variation observed for the Omicron BA.1, Omicron BA.5 and the joint removal of both Omicron BA.2 variants (fig. S18). Similarly, variant positions were stable to the removal of serum groups, albeit with slightly larger variation when removing the BA.1 and BA.5 convalescent sera.

Finally, we investigated how well the map is able to predict missing titrations. To this end, we performed 500 cross-validation repeats, where 10% of the titers were removed at random, the map re-optimised, and the missing titers predicted from the antigenic map. The mean difference between the predicted and measured titer was 0.44, with a standard deviation of 1.36 when assuming a mean of 0. This is higher than the difference between fitted and measured titers (fig. S15, mean = 0.14, standard deviation = 0.73 when assuming a mean of 0 and only considering detectable titers). We visualized the difference between predicted and measured titers split by variant and serum group, and found that no variant is predicted to consistently have higher residuals in all serum groups.

### Supplementary figures

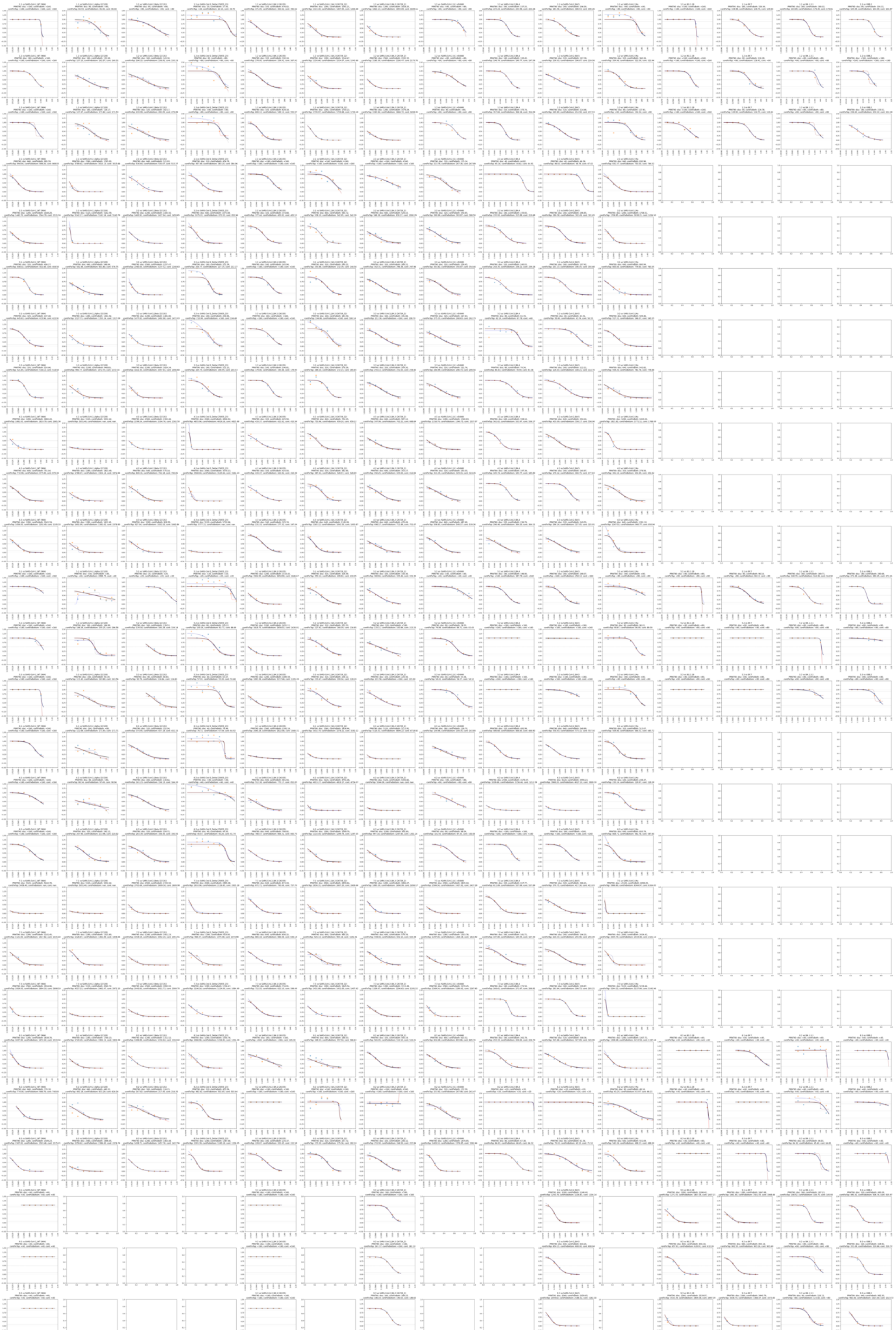

**Figure S1: Neutralisation curves.** Each sub-plot shows the neutralization curve for a variant titrated against a serum, as specified in the panel title. The y-axis shows the fraction of infectivity remaining, the x-axis shows the dilution. Blue and orange shapes show the fraction of infectivity remaining from the two runs that were done. Titer curves were fit with the neutcurve package (*1*). The black curve shows the titer curve inferred when constraining the lower end of the neutralization curve at zero and the upper end at one. The red curve shows the titer curve inferred when constraining the upper end of the neutralization curve at one. The blue curve shows the titer curve inferred when constraining the bottom end of the neutralization curve at 0, and the gray curve shows the titer curve inferred when not constraining the neutralization curve.

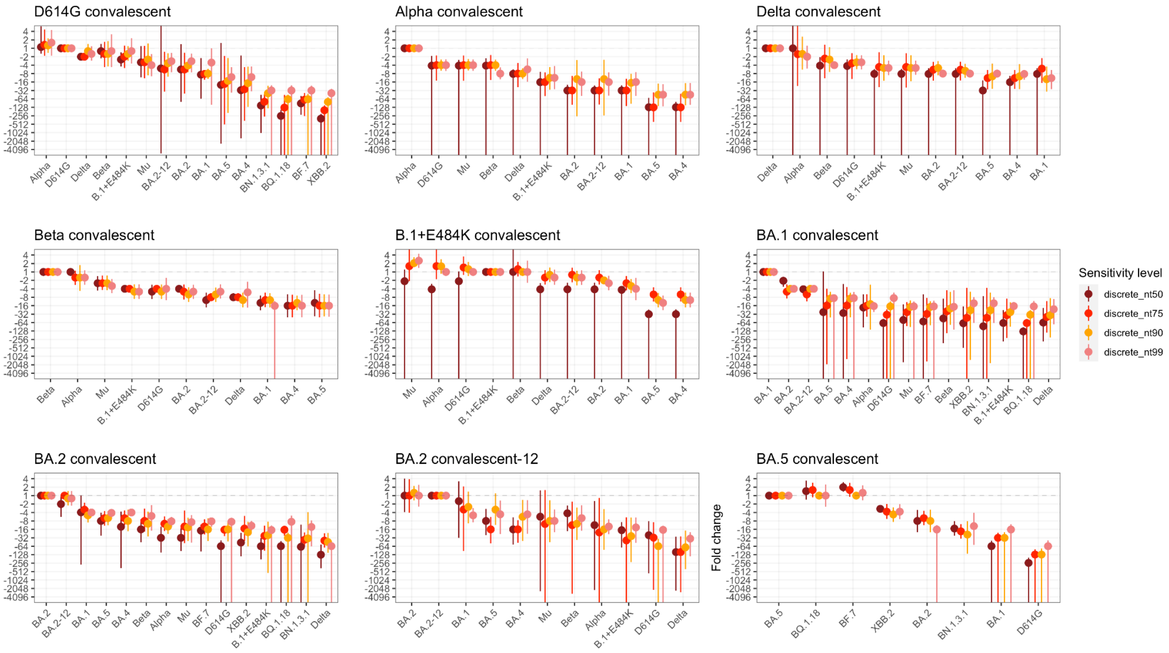

**Figure S2: Fold change measured when inferring discrete titers with different sensitivity levels.** Larger fold changes with wide confidence intervals result when titers are non-detectable. Fold changes are largely comparable between sensitivity levels. Dots show the estimated fold change, with lines showing the 95% highest posterior density interval.

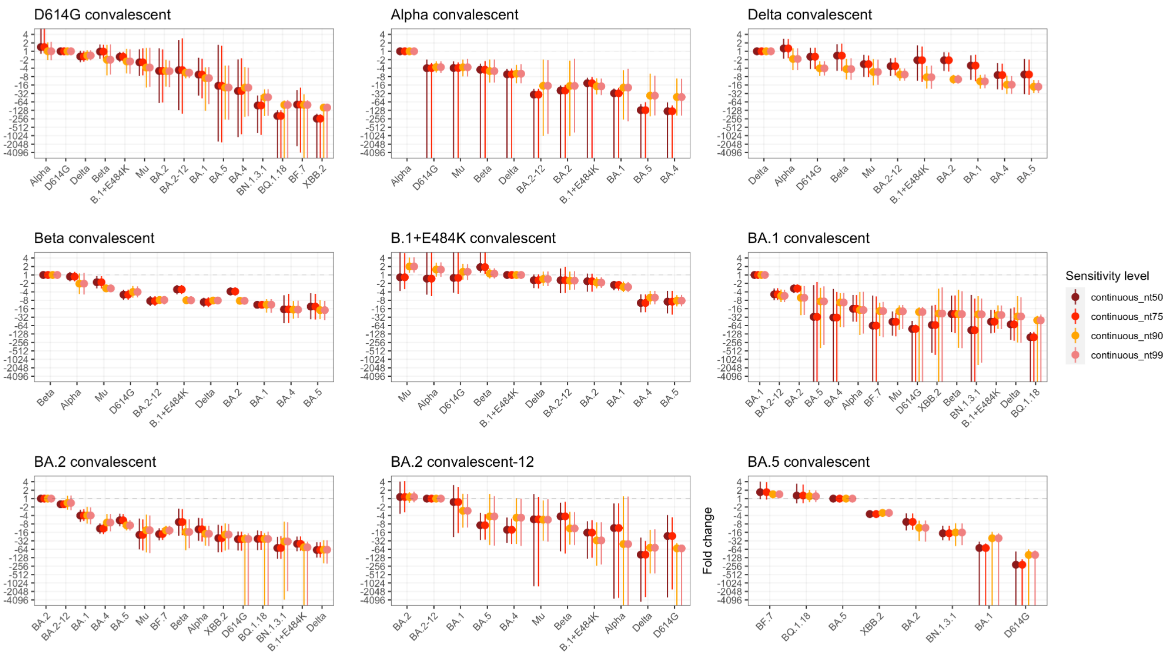

**Figure S3: Fold change measured when inferring continuous titers with different sensitivity levels.** Larger fold changes with wide confidence intervals result when titers are non-detectable. Fold changes are largely comparable between sensitivity levels. Continuous titers were inferred by constraining the neutralization curve at 0 and 1. Dots show the estimated fold change, with lines showing the 95% highest posterior density interval.

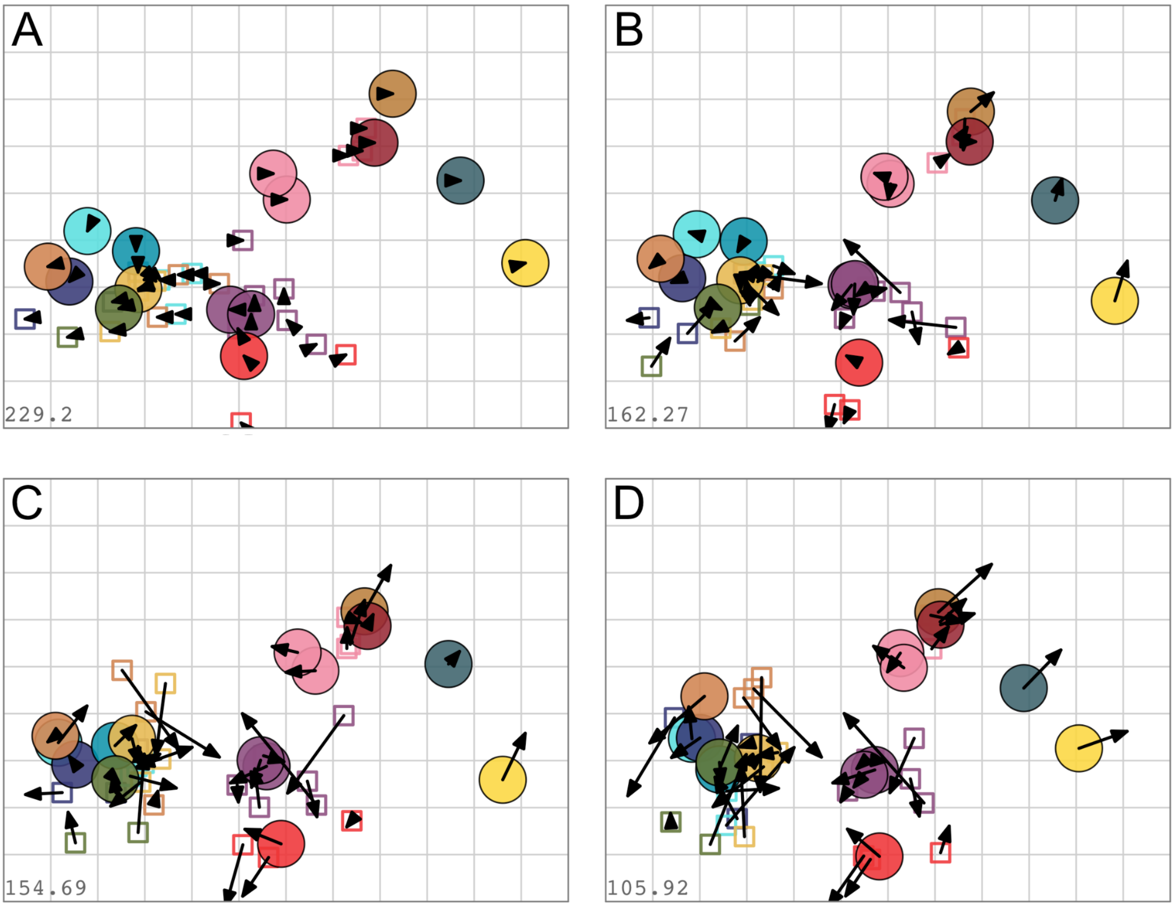

**Figure S4: Antigenic maps made from discrete titers inferred with different sensitivity levels.** Antigenic maps were optimized in two dimensions, with 500 optimisation and the minimum column basis parameter set to “none”. Maps were inferred using a dilution_stepsize of 1. Arrows point to the position of the same variant and serum in the NT50 map shown in panel A. A) NT50, B) NT75, C) NT90, D) NT99.

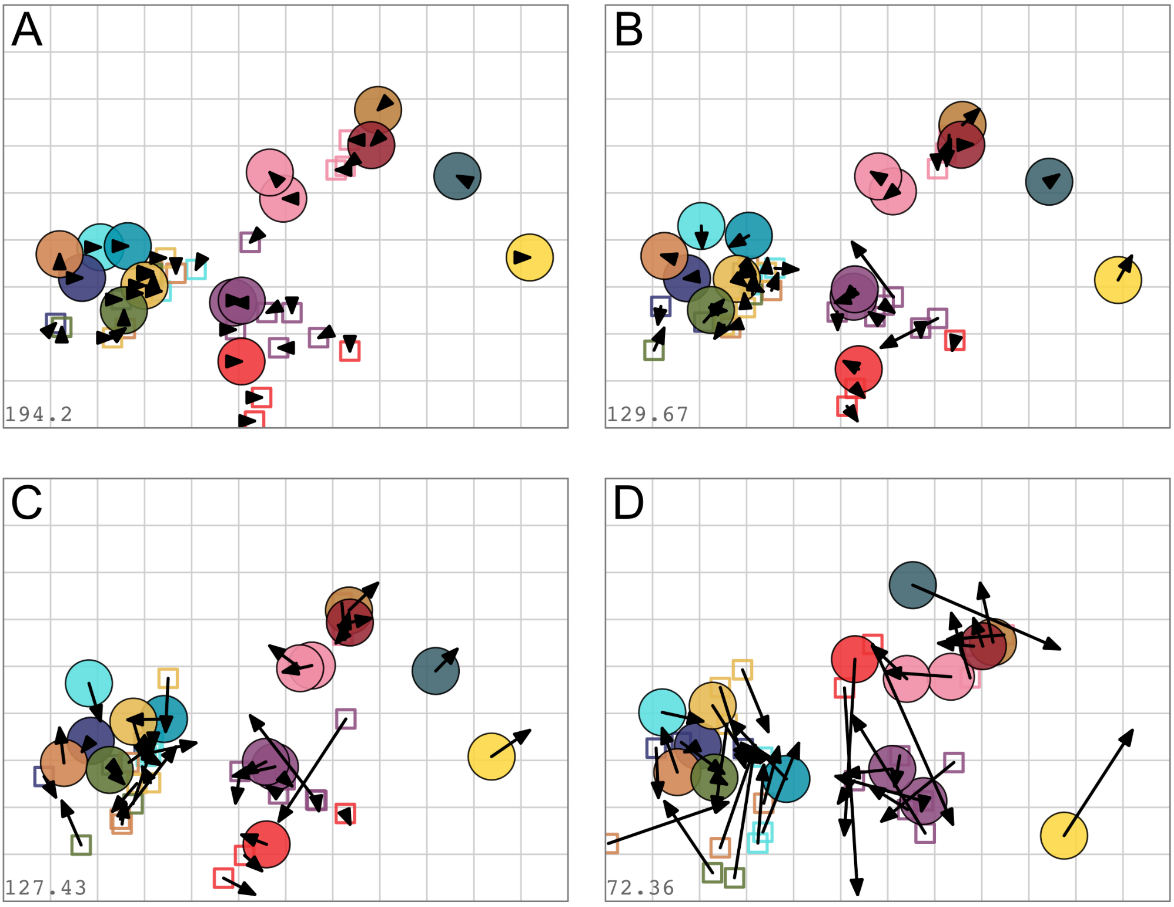

**Figure S5: Antigenic maps made from continuous titers inferred with different sensitivity levels.** Antigenic maps were optimized in two dimensions, with 500 optimisation and the minimum column basis parameter set to “none”. Maps were inferred using a dilution_stepsize of 0. Arrows point to the position of the same variant and serum in the NT50 map shown in panel A. A) NT50, B) NT75, C) NT90, D) NT99. Continuous titers were inferred by constraining the neutralization curve at 0 and 1.

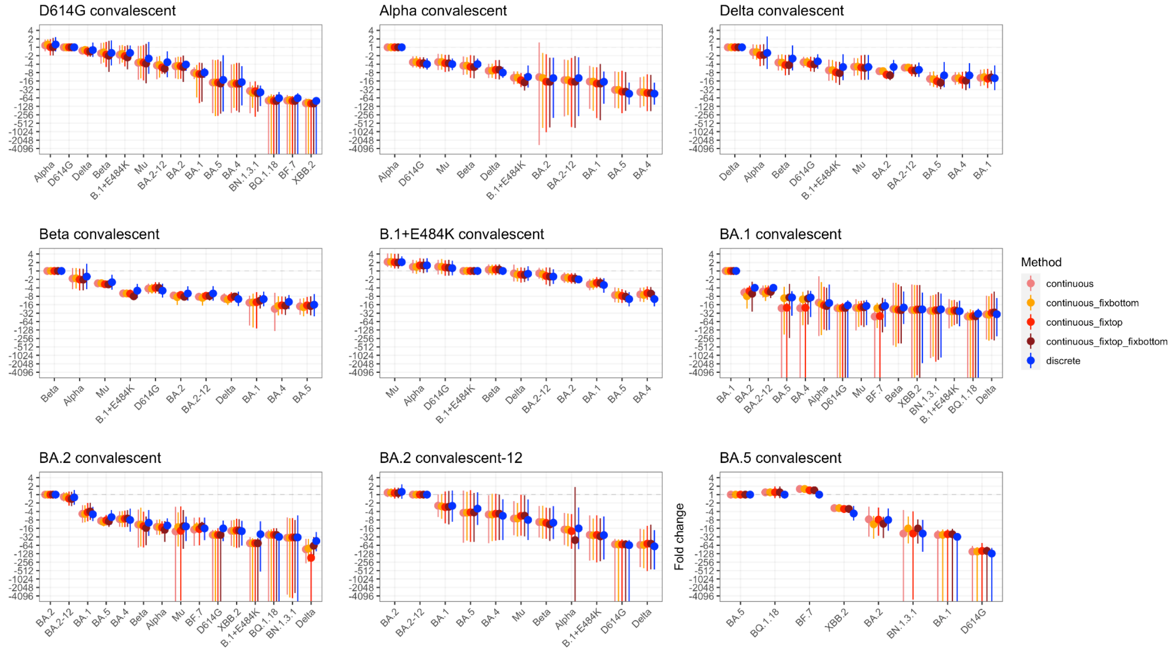

**Figure S6: Fold change measured when inferring titers using either continuous or discrete methods at NT90.** Continuous titers were inferred fixing the bottom and top of the neutralization curve at zero and one, respectively (continuous_fixtop_fixbottom), only fixing the top of the neutralization curve (continuous_fixtop), only fixing the bottom of the neutralization curve (continuous_fixbottom), not fixing the neutralization curve (continuous), and using discrete titers (discrete). Fold changes are similar between different methods. Dots show the estimated fold change, with lines showing the 95% highest posterior density interval.

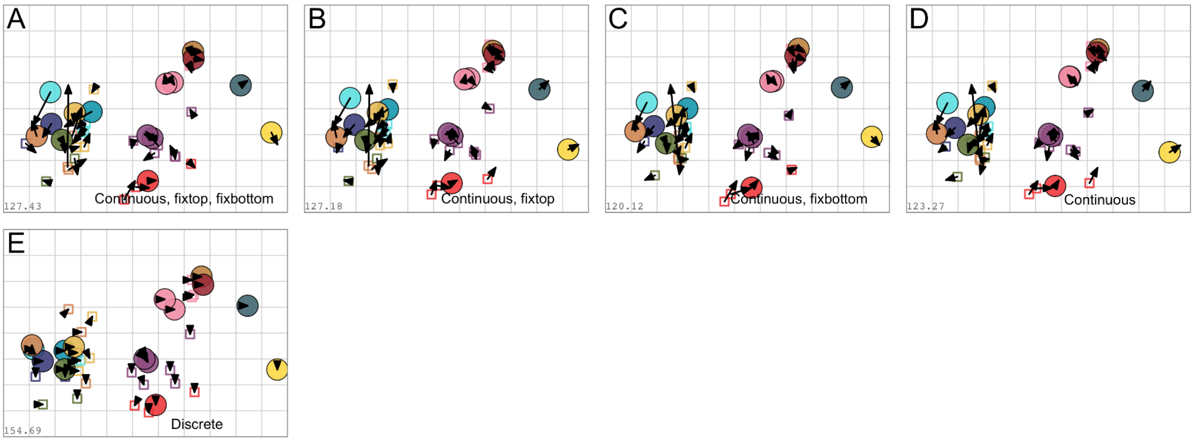

**Figure S7: Comparison of discrete and continuous NT90 maps.** In each map, the arrows point to the position of the variant and serum in the antigenic map made from discrete titers, shown in panel E. A) Map made from continuous titers, fixing the bottom and top of the neutralization curve at zero and 1, respectively. B) Map made from continuous titers, fixing the top of the neutralization curve at one. C) Map made from continuous titers, fixing the bottom of the neutralization curve at one. D) Map made from continuous titers, without fixing the neutralization curve. E) Map made from discrete titers.

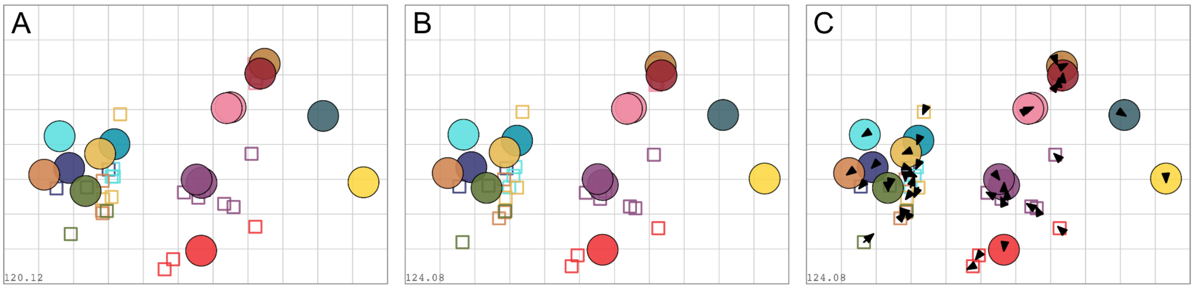

**Figure S8: Comparison of the PRNT90 fixbottom map and the PRNT90 fixbottom map with a subset of titers manually adapted.** A) PRNT90 map inferred from titers calculated while fixing the bottom of the neutralization curve at zero ('fixbottom map'). B) PRNT90 map inferred from titers calculated while fixing the bottom of the neutralization curve at zero, with a subset of titers manually adapted. C) PRNT90 fixbottom map with a subset of titers manually adapted with arrows pointing to the positions of sera and variants in the non-adapted map. The adapted titers were: 2.1 vs Alpha, 2.2 vs BA.2_2, 3.2 vs BA.2_12, 3.3 vs Alpha, 3.3 vs Beta, 4.1 vs Alpha, 4.2 vs Delta, 4.3 vs Alpha, 4.3 vs Delta, 5.1 vs Alpha, 5.1 vs BA.2_12, 5.2 vs Alpha, 5.2 vs BA.2_2, 5.3 vs BA.2_2, 6.1 vs BA.2_2, 6.2 vs BA.2_2, 7.1 vs D614G, 7.1 vs Alpha, 7.1 vs Delta, 7.1 vs E484K, 7.1 vs BA.2_12, 7.1 vs Mu, 7.2 vs Mu, 7.3 vs Alpha, 7.3 vs Mu, 8.1 vs BA.2_12, 9.1 vs BF.7, 9.3 vs BQ.1.18. Adaptations were made as shown in Table S2.

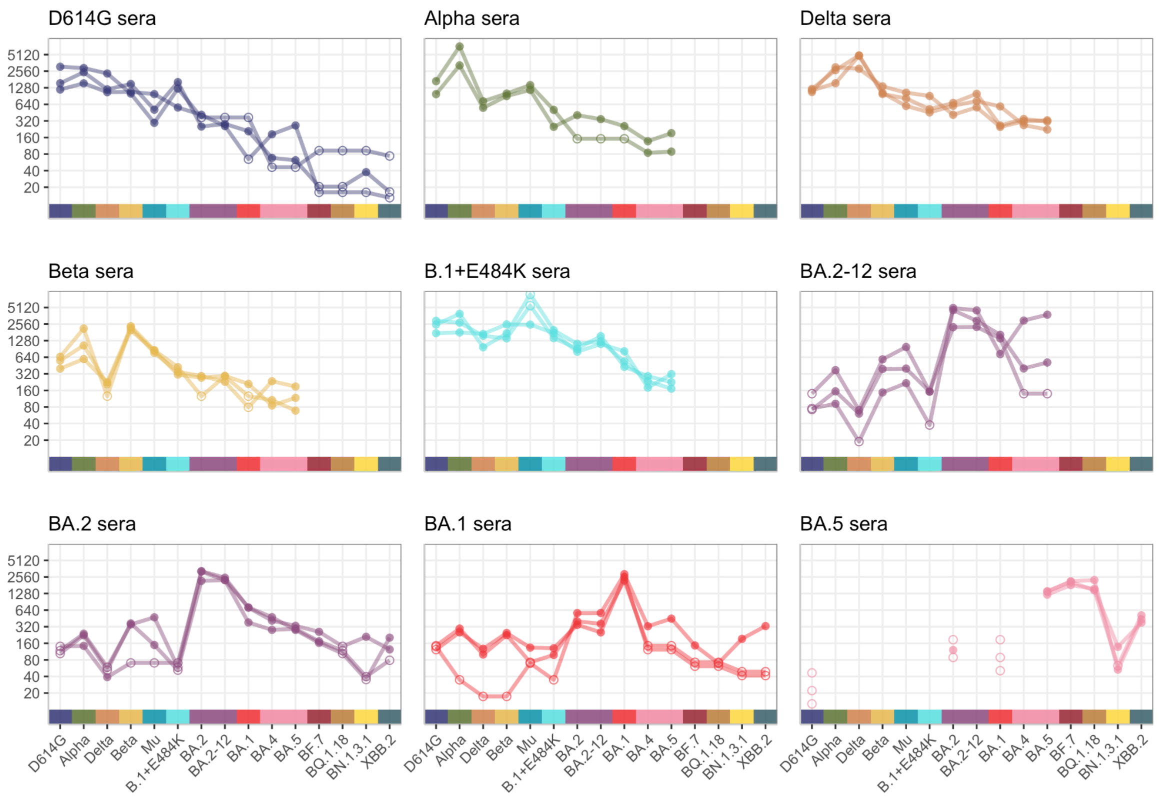

**Figure S9: Neutralizing titers of 15 live virus isolates titrated against sera from hamsters twice infected with D614G, Alpha, Beta, Delta, Mu, B.1+E484K, Omicron BA.1, Omicron BA.2 (two isolates), or Omicron BA.5, after adjusting for differences in individual serum magnitude.** Variants that were titrated are the pre-Omicron variants D614G, Alpha, Beta, Delta, Mu, B.1+E484K, first generation Omicron variants BA.1, BA.2, BA.4, BA.5, second generation Omicron variants BF.7, and BQ.1.18, third generation Omicron variant BN.1.3.1, and the recombinant Omicron XBB.2. Detectable titers are shown as filled circles, non-detectable titers are shown as empty circles. Following (*2*), to account for differences in individual serum magnitude (where one individual may have consistently higher or lower titers), each titer of variant v against serum s was given by $t_{v,s} = s_{v,S} + r_s + v_v + e$ where $s_{v,S}$ is the average logtiter of variant v against serum group S, $r_s$ is the individual serum magnitude effect, $v_v$ is the antigen reactivity effect (accounting for a particular variant having high titers against all sera) and e is independently and normally distributed log titer noise. The ‘cmdstanr’ R package (*4*) was used as an interface to the Stan (*5*) optimisation procedures for fitting the model. The estimated individual serum magnitude effects were subtracted from each titer to adjust for individual serum magnitude differences. Reactivity patterns among the hamsters infected with the same isolate were largely uniform with the exception of one B.1+E484K serum with a lower titer against Mu, one D614G serum and one BA.2 serum both with increased reactivity against BA.4 and BA.5, and one BA.1 serum with higher titers against the Omicron variants BA.4, BA.5, BF.7, BN.1.3.1, and XBB.2, and lower titers against Alpha, Delta, and Beta.

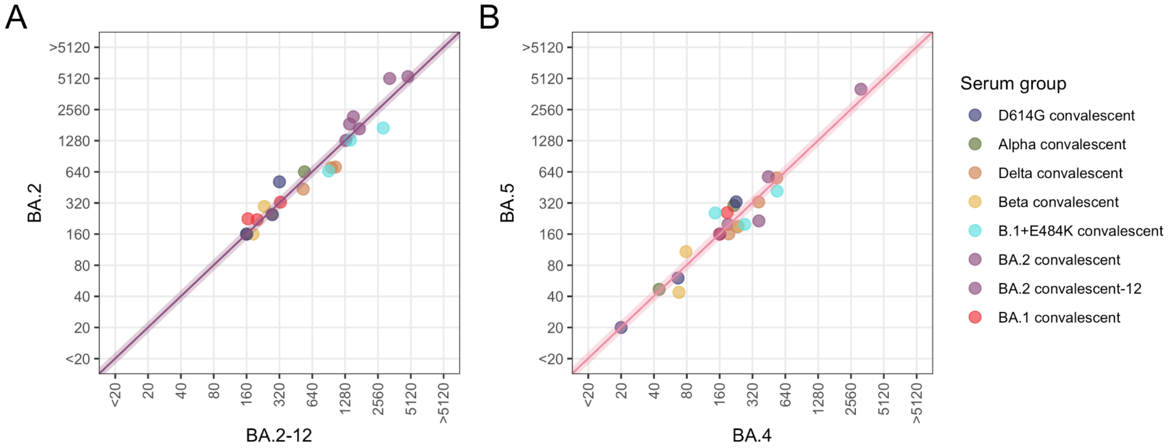

**Figure S10: Pairwise comparison of titers of two different variants.** A) Comparison of the titers from the two different BA.2 isolates. The mean fold difference between the two variants is 1 (95% highest posterior density interval: -1.14, 1.13). B) Comparison of the titers from the BA.4 and BA.5 variants. The mean fold difference between the two variants is -1 (95% highest posterior density interval: -1.16, 1.15). Dots are coloured by serum group. The intercept of the coloured line shows the mean fold difference between the titers from the two variants, the slope is equal to 1. The shaded area shows the 95% posterior density interval of the fold difference.

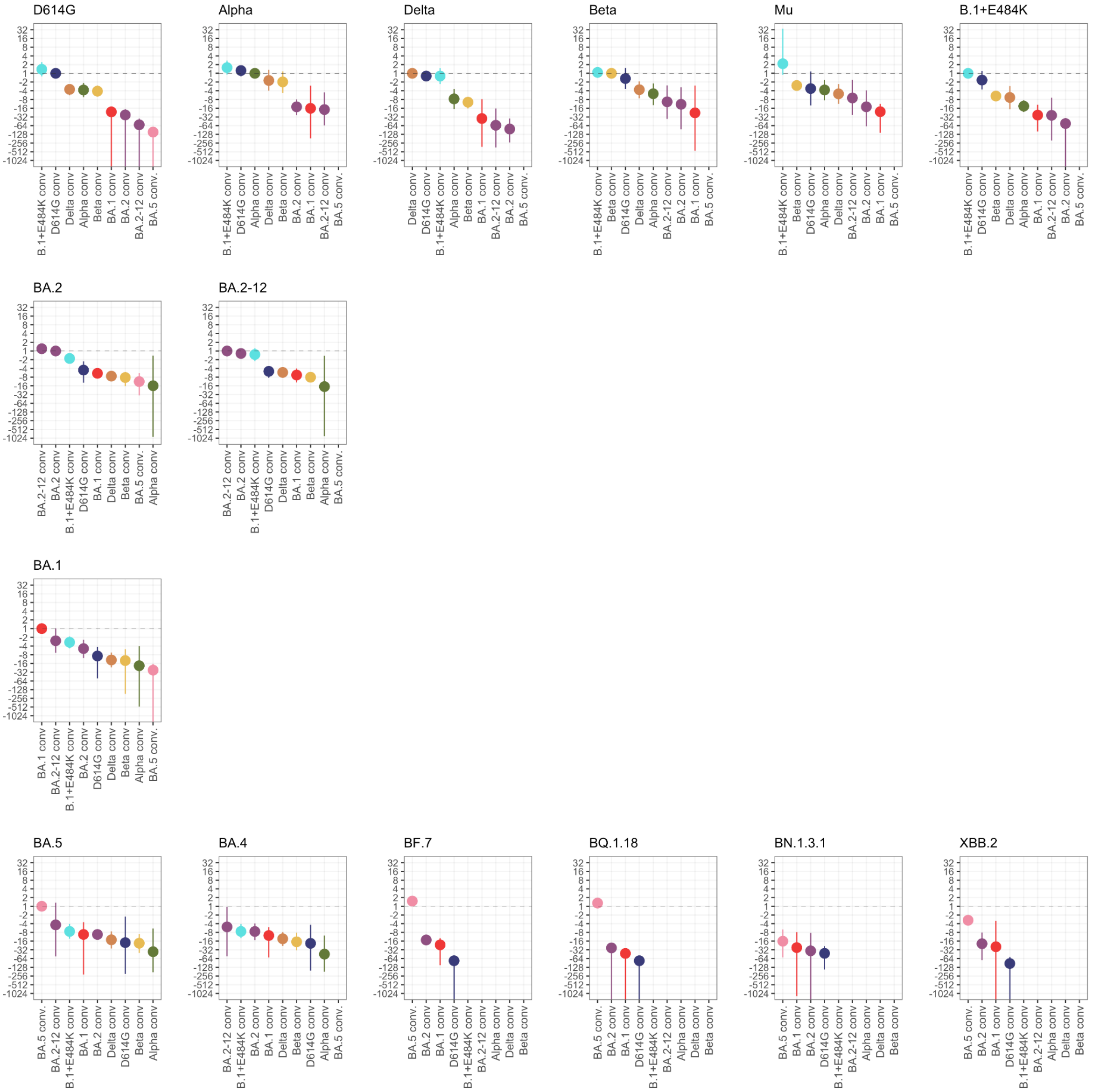

**Figure S11: Fold change compared to the homologous variant aggregated by variant.** The serum groups on the x-axis are ordered by decreasing fold change for that variant. Dots indicate the estimated mean fold change, the bars the 95% highest posterior density intervals.

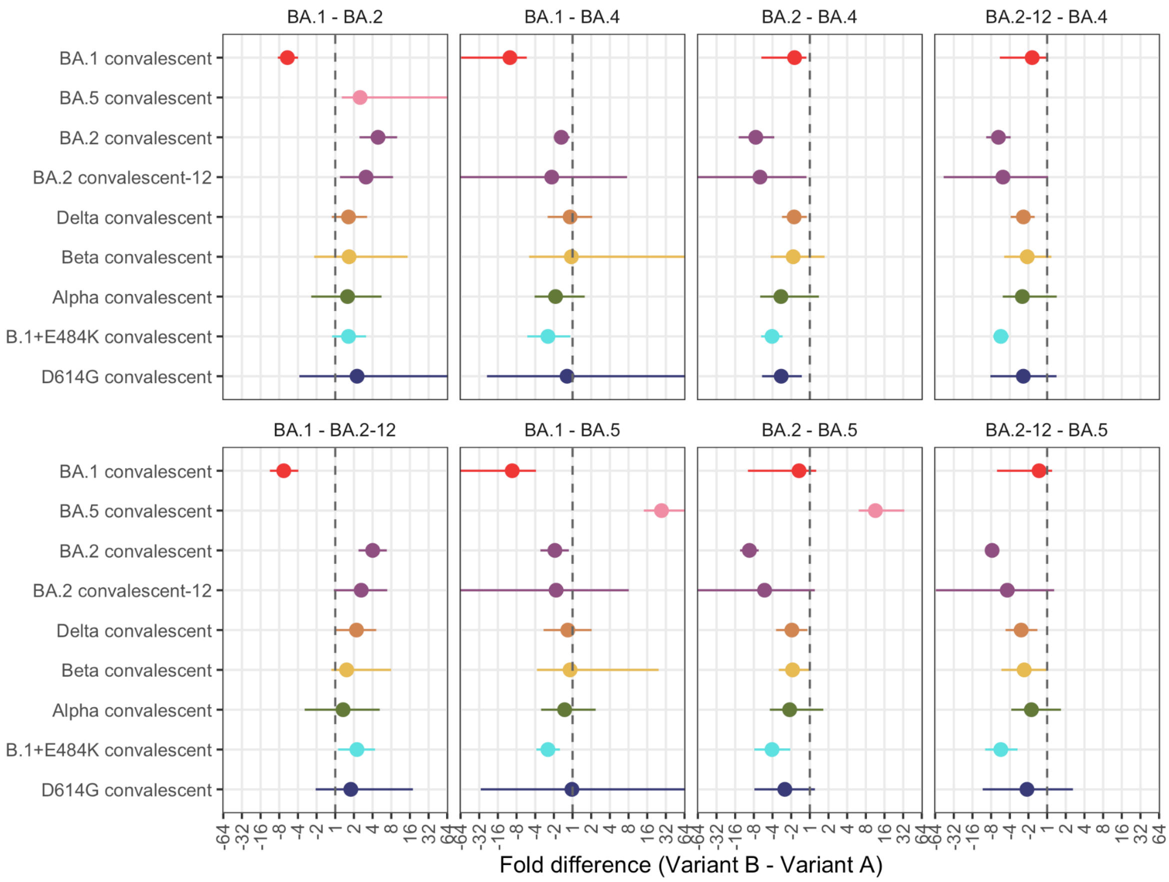

**Figure S12: Fold change between Omicron variants as measured by different serum groups.** Dots indicate the estimated mean fold change, the bars the 95% highest posterior density intervals.

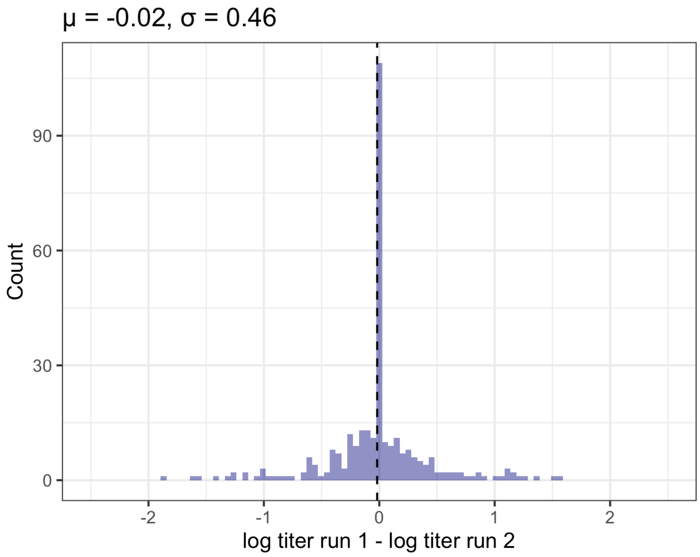

**Figure S13: Repeat variation.** Each titer was measured in duplicate. The figure shows the difference in $log_2$ titer between the first and the second repeat. Non-detectable titers were set to limit of detection - 1 on the $log_2$ scale. The mean difference between repeats was 0.02, with a standard deviation of 0.46. Assuming a mean of 0 to account for the systematic bias of titrations between repeats, the standard deviation is 0.46 on the $log_2$ scale. Accounting for the presence of measurement error in both the first and second titration the standard deviation of noise per measurement is $\sqrt{0.46^2/2} = 0.32$.

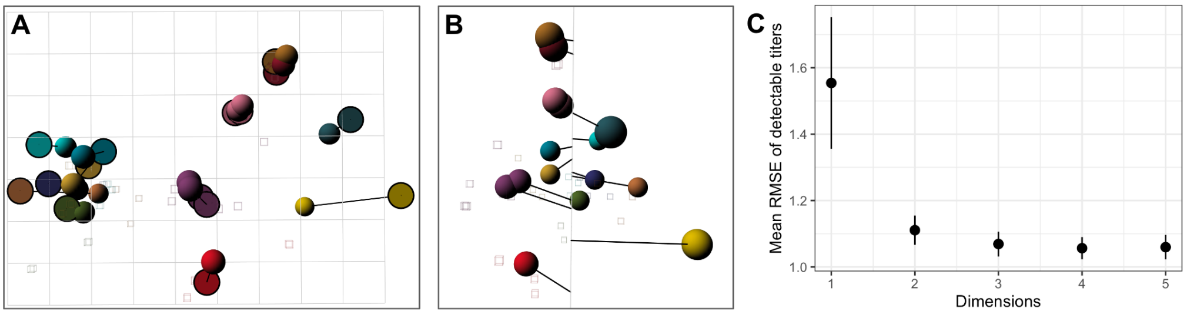

**Figure S14: Testing map dimensionality.** A and B) The map optimized in three dimensions, shown from the side (A) and from the front (B). The black lines point to the position of the variants in the two-dimensional map shown in Figure 3. C) Dimensionality test, where 1000 repeats were performed with 10% of titers excluded at random and the map optimized in one to five dimensions. The plot shows the root mean squared error and standard error of the excluded titers between the titers estimated from the map and the known titers.

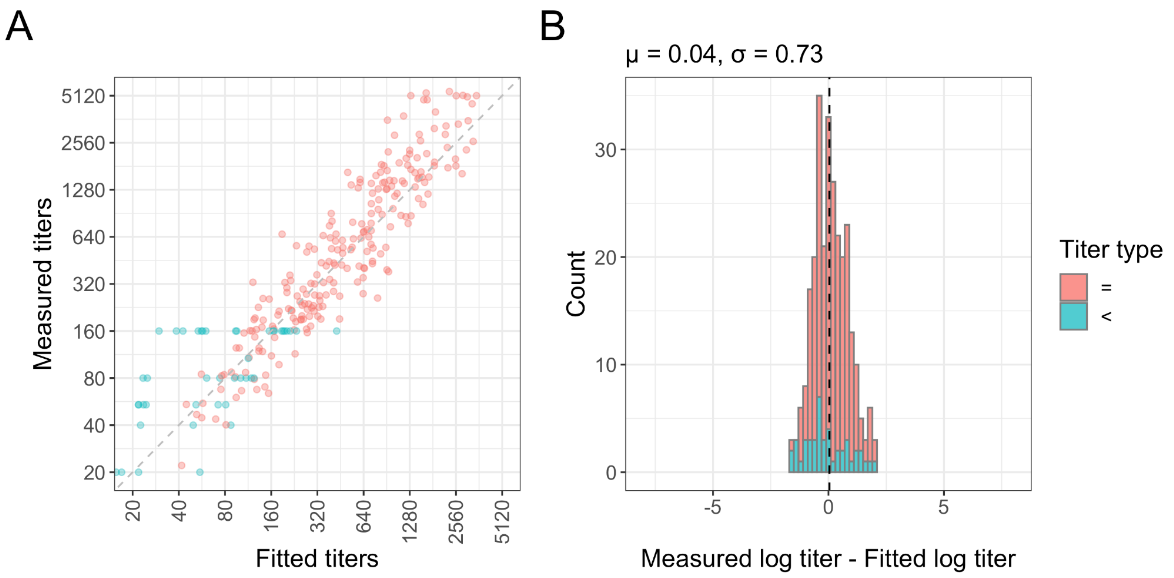

**Figure S15: Comparison of measured $log_2$ titers and fitted $log_2$ titers as inferred from the antigenic map.** Detectable measured titers are shown in red, non-detectable measured titers in blue. A) Scatter plot of measured and fitted $log_2$ titers. Fitted $log_2$ titers were inferred from the distances in the antigenic map. B) Histogram of the difference between the measured and the fitted $log_2$ titers. When measured titers were non-detectable, the residuals were inferred as described in (*2*). Variant and serum group pairs where all titers were non-detectable were excluded (BA.2 convalescent vs D614G and B.1+E484K, BA.1 convalescent vs D614G, BA.2 convalescent-12 vs D614G). Measured titers are on average 0.04 higher than the fitted titers (with a standard deviation of 0.73), with the mean indicated by the dashed black line. Assuming a mean of 0 to account for the systematic bias of titrations between repeats, the standard deviation is 0.73 on the $log_2$ scale. This is higher than the standard deviation of the variation between repeats, however the standard deviation between repeats may be lower, since the repeats were not independent. When only including detectable titers, the mean difference between measured and fitted $log_2$ titers was 0.14 with a standard deviation of 0.73 when assuming a mean of 0.

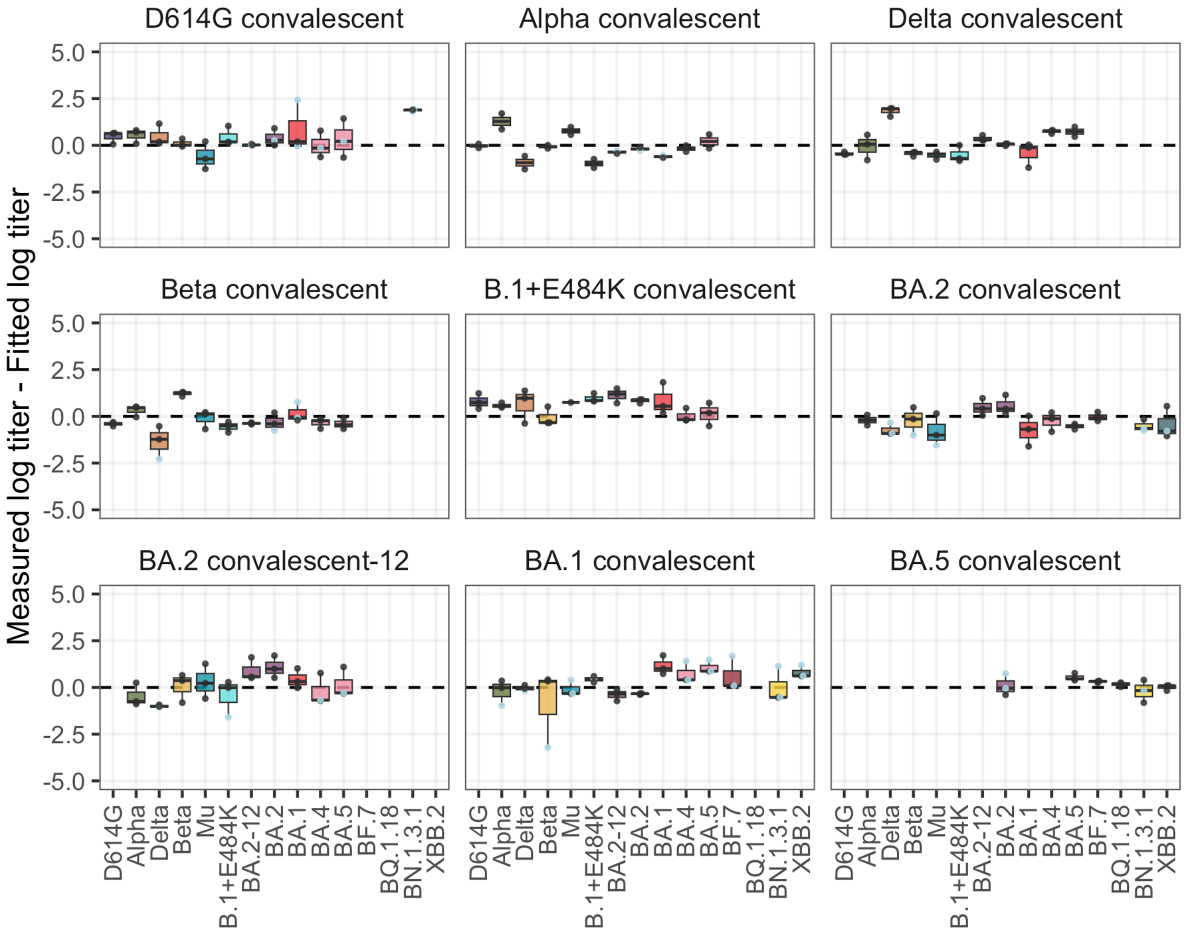

**Figure S16: The difference between the measured $\log_2$ titer and the fitted $\log_2$ titer as inferred from the antigenic map, split by serum group and variant.** Residuals for detectable titers are shown in black, residuals for non-detectable titers in blue. Variant and serum group pairs where all titers were non-detectable were excluded (BA.2 convalescent vs D614G and B.1+E484K, BA.1 convalescent vs D614G, BA.2 convalescent-12 vs D614G). The boxplot indicates the median and 25$^{th}$ and 75$^{th}$ percentile.

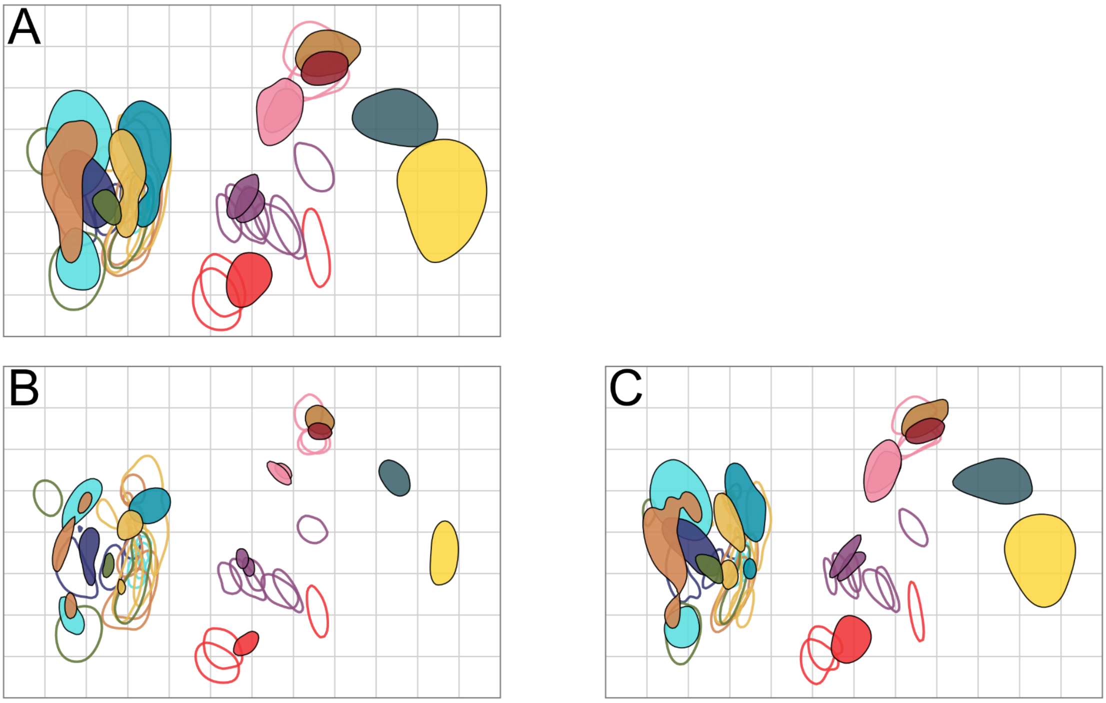

**Figure S17: Assessing sensitivity of the antigenic map to measurement error.** 1000 bootstrap repeats were performed where normally distributed noise was added to each titer and / or to the titers of each variant. A) Noise added to titers and variant reactivity. B) Noise added to titers only. C) Noise added to variants only. Normally distributed noise added to the titers had a standard deviation of 0.32, following the standard deviation of the measurement error estimated in figure S13. Normally distributed noise added on a per-variant basis had a standard deviation of 0.4. The coloured regions indicate the area where 68% (1 standard deviation) of the positional variation of each variant (filled blobs) and serum (empty blobs) is captured.

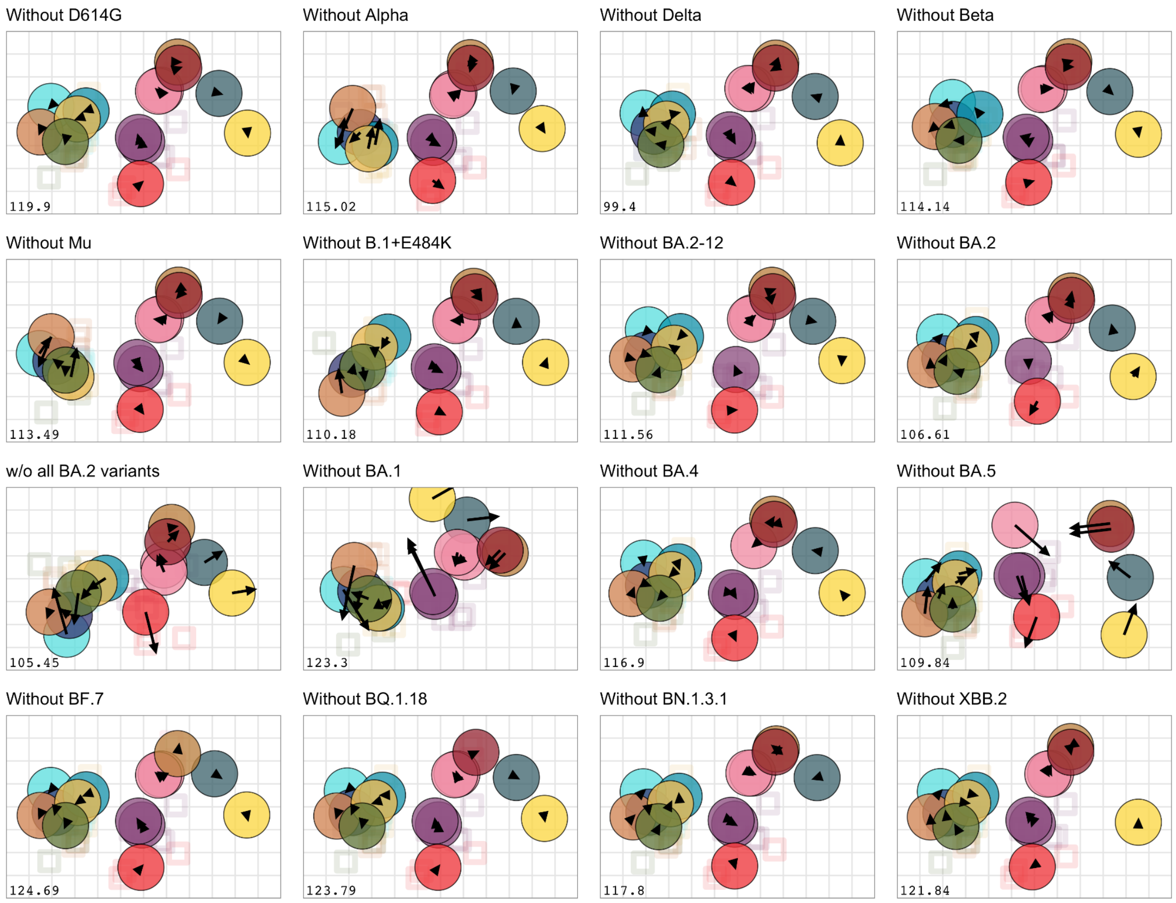

**Figure S18: Robustness of the antigenic map to removing variants.** Each variant was removed in turn, as indicated in the panel title, and the map re-optimised. The arrows point to the position of the variants in the full map, shown in Figure 3.

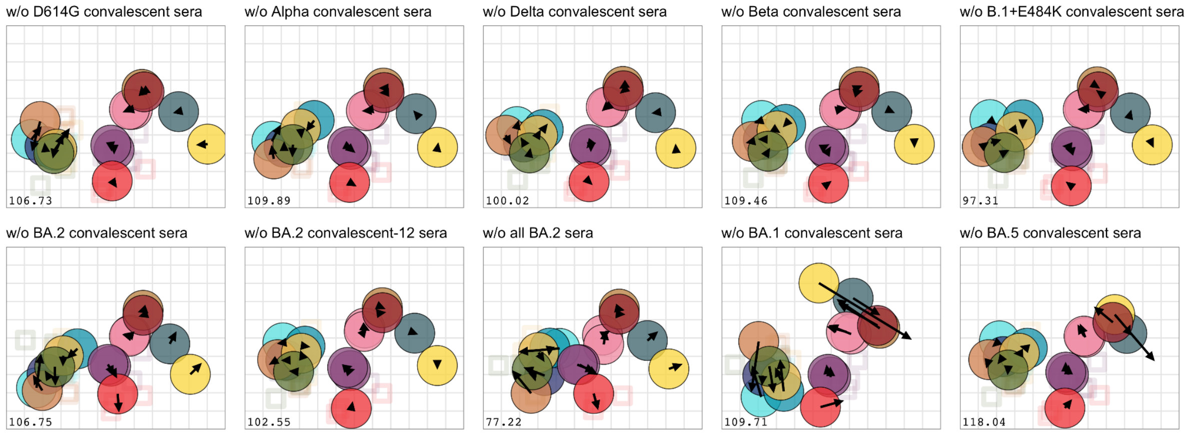

**Figure S19: Robustness of the antigenic map to removing sera.** Each group of sera was removed in turn, as indicated in the panel title, and the map re-optimised. The arrows point to the position of the variant in the map with all serum groups shown in Figure 3.

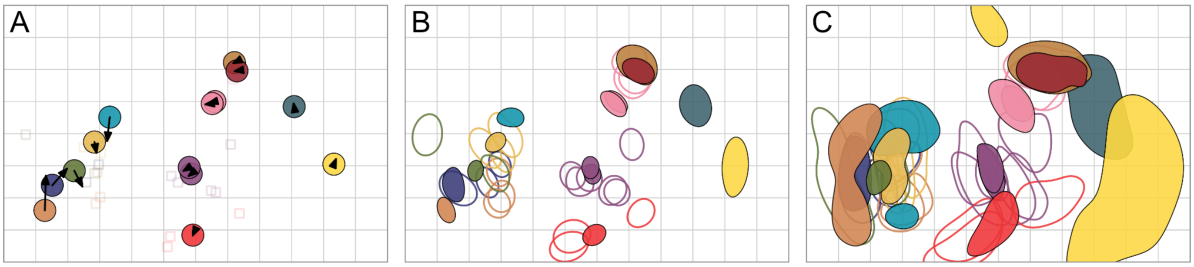

**Figure S20: Antigenic map made without the B.1+E484K variant and sera.** A) Antigenic map inferred without the B.1+E484K variant and sera, with arrows pointing to the positions of the variants in the map with the B.1+E484K variant and sera (as in Figure 3). B) Antigenic map with triangulation blobs. The blobs show the area that each variant (filled blob) or serum (empty blob) could take up without increasing the stress of the map by 1. C) Antigenic map with resampling bootstrap blobs. An antigenic map was inferred 500 times, and each time titers were sampled with replacement. The area of the blobs indicate the area that each variant (filled blob) or serum (empty blob) occupy in 68% of the repeats. The coloring of the variants and sera is as in Figure 3.

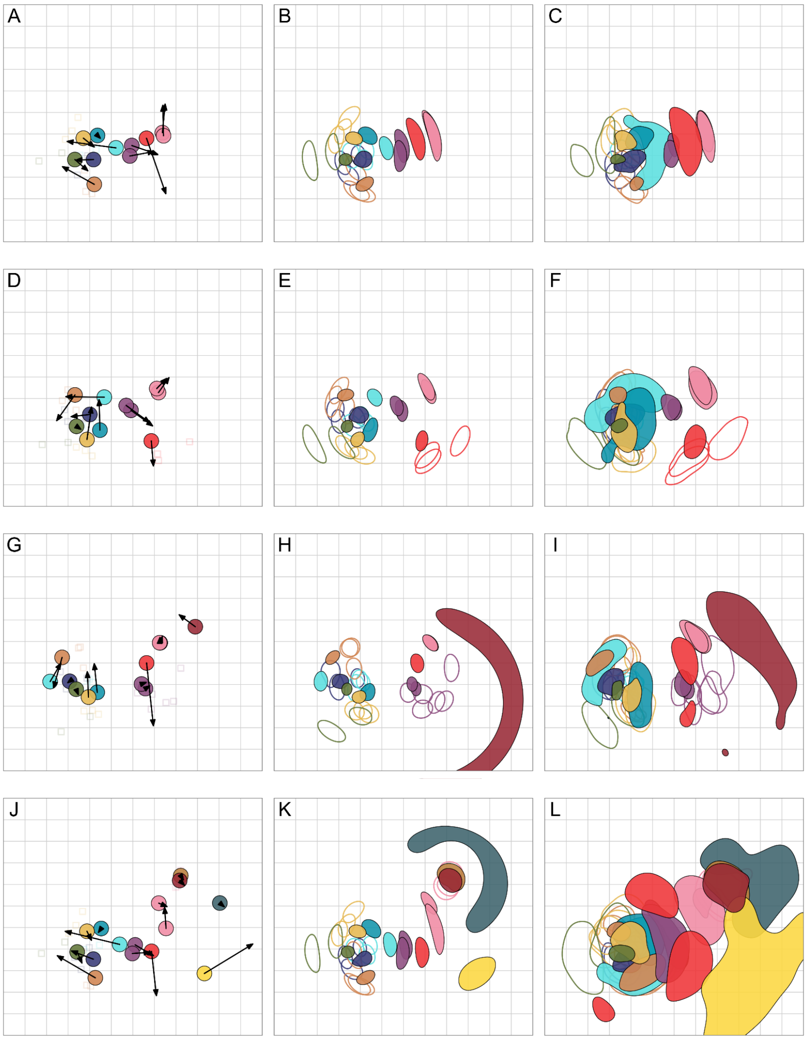

**Figure S21: Antigenic map made with different combinations of Omicron sera.** The left column always shows the antigenic map inferred without a subset of Omicron sera, with the arrows pointing to the positions of the variants in the full map (as in Figure 3). The middle column shows the antigenic map inferred without a subset of Omicron sera with triangulation blobs. The blobs show the area that each variant (filled blob) or serum (empty blob) could take up without increasing the stress of the map by more than 1. The right column shows the antigenic map inferred without a subset of Omicron sera with resampling bootstrap blobs. An antigenic map was inferred 500 times, and each time titers were sampled with replacement. The area of the blobs indicate the area that each variant (filled blob) or serum (empty blob) occupy in 68% of the repeats. The coloring of the variants and sera is as in Figure 3. A-C) Antigenic maps without any Omicron sera (the BF.7, BQ.1.18, XBB.2, and BN.1.3.1 variants cannot be placed on the map due to insufficient detectable titers). D-F) Antigenic maps without BA.2 and BA.5 convalescent sera (the BF.7, BQ.1.18, XBB.2, and BN.1.3.1 variants cannot be placed on the map due to insufficient detectable titers). G-I) Antigenic maps without BA.1 and BA.5 convalescent sera. J-L) Antigenic maps without BA.1 and BA.2 convalescent sera.

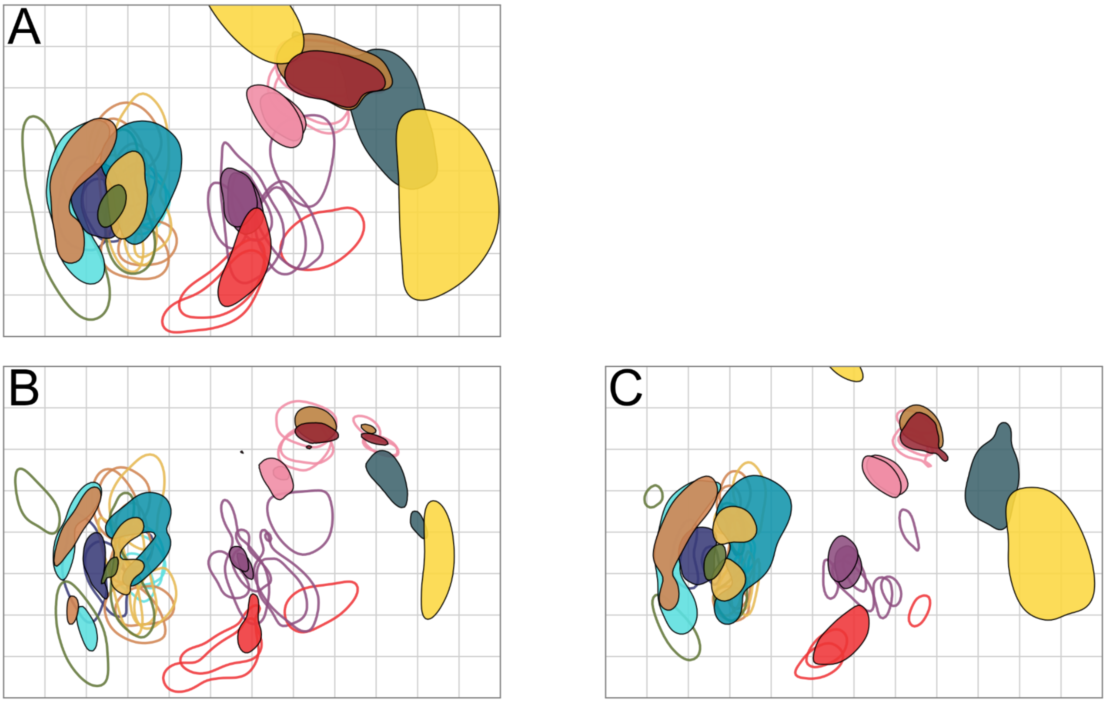

**Figure S22: Robustness of the antigenic map to missing titers.** 1000 bootstrap repeats were performed, where a random subset of titers was sampled with replacement, and the map re-optimised. A) Sera and variants were re-sampled. B) Variants were resampled. C) Sera were resampled. The coloured regions indicate the area where 68% (1 standard deviation) of the positional variation of each variant (filled blobs) and serum (empty blobs) is captured.

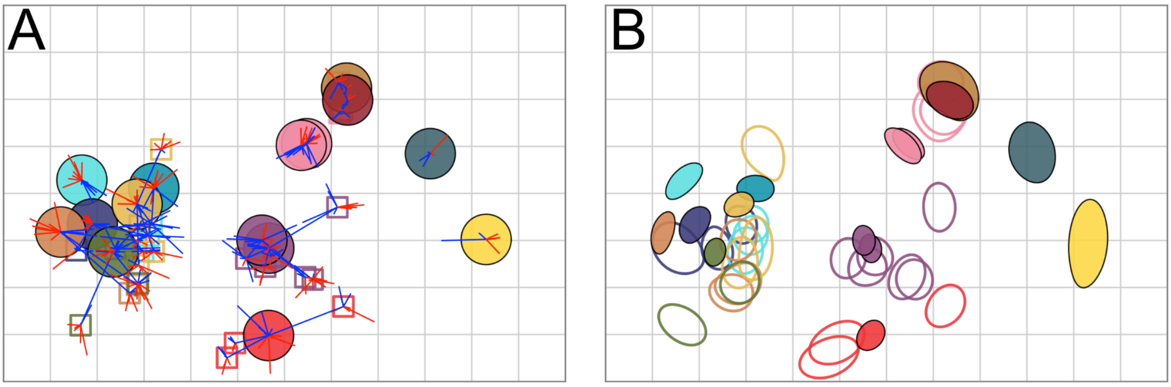

**Figure S23: Uncertainty of the position of variants and sera.** A) Antigenic map with error lines. For each variant/serum pair a blue or red line is plotted showing the target distance between the variant and serum. A blue line indicates a target distance smaller, a red line a distance larger than the one shown in the map. B) Antigenic map with triangulation blobs. The blobs show the area that each variant (filled blob) or serum (empty blob) could take up without increasing the stress of the map by 1.

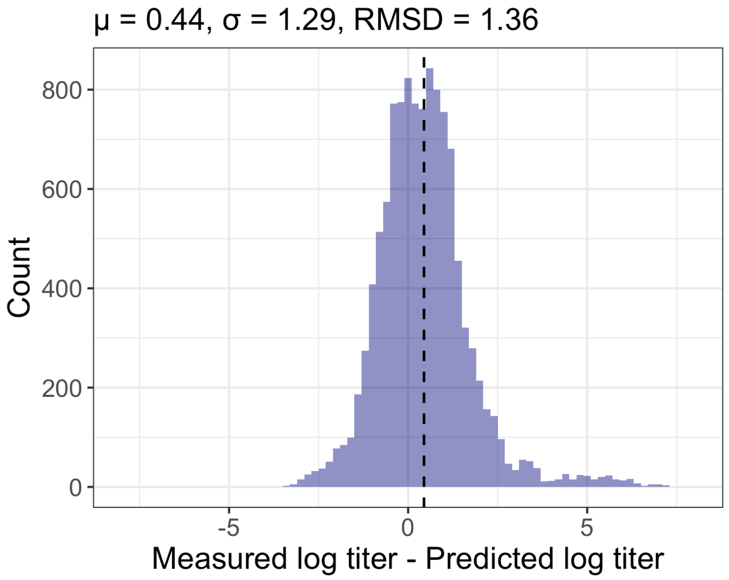

**Figure S24: Assessing predictive power of the antigenic map.** 500 repeats were performed where the map was re-optimized leaving out 10% of the data at random. The histogram shows the difference between the measured $\log_2$ titer and the predicted $\log_2$ titer as inferred from the map with 10% of the data removed. Only detectable titers were considered. The mean difference between measured and predicted detectable titers on the $\log_2$ scale was 0.44, with a standard deviation of 1.29. Assuming a mean of 0, the standard deviation is 1.36.

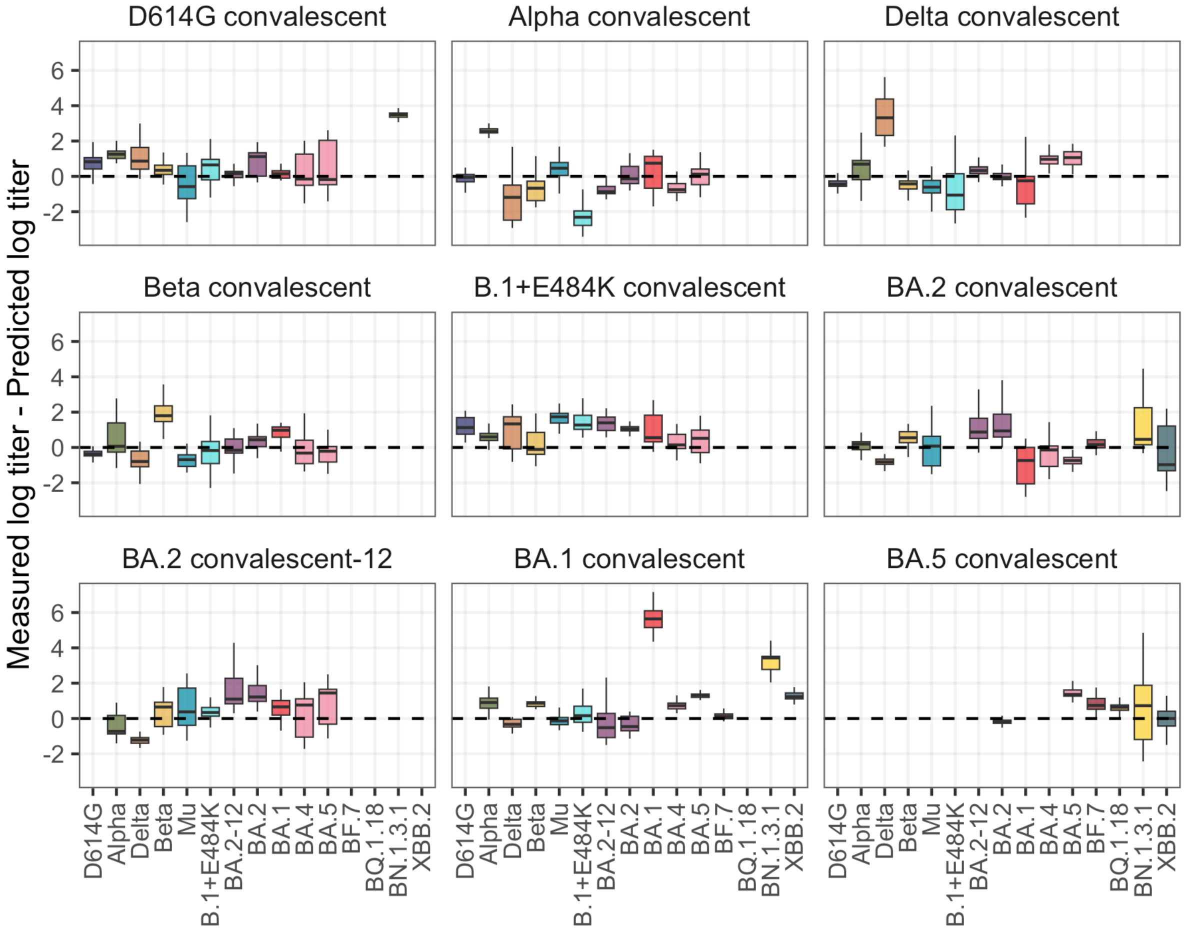

**Figure S25: Assessing predictive power of the antigenic map, split by serum group and variant.** The difference between measured and predicted $log_2$ titers was inferred as described in the legend of figure S25. Only detectable titers are considered. The boxplot shows the median and 25th and 75th percentile.

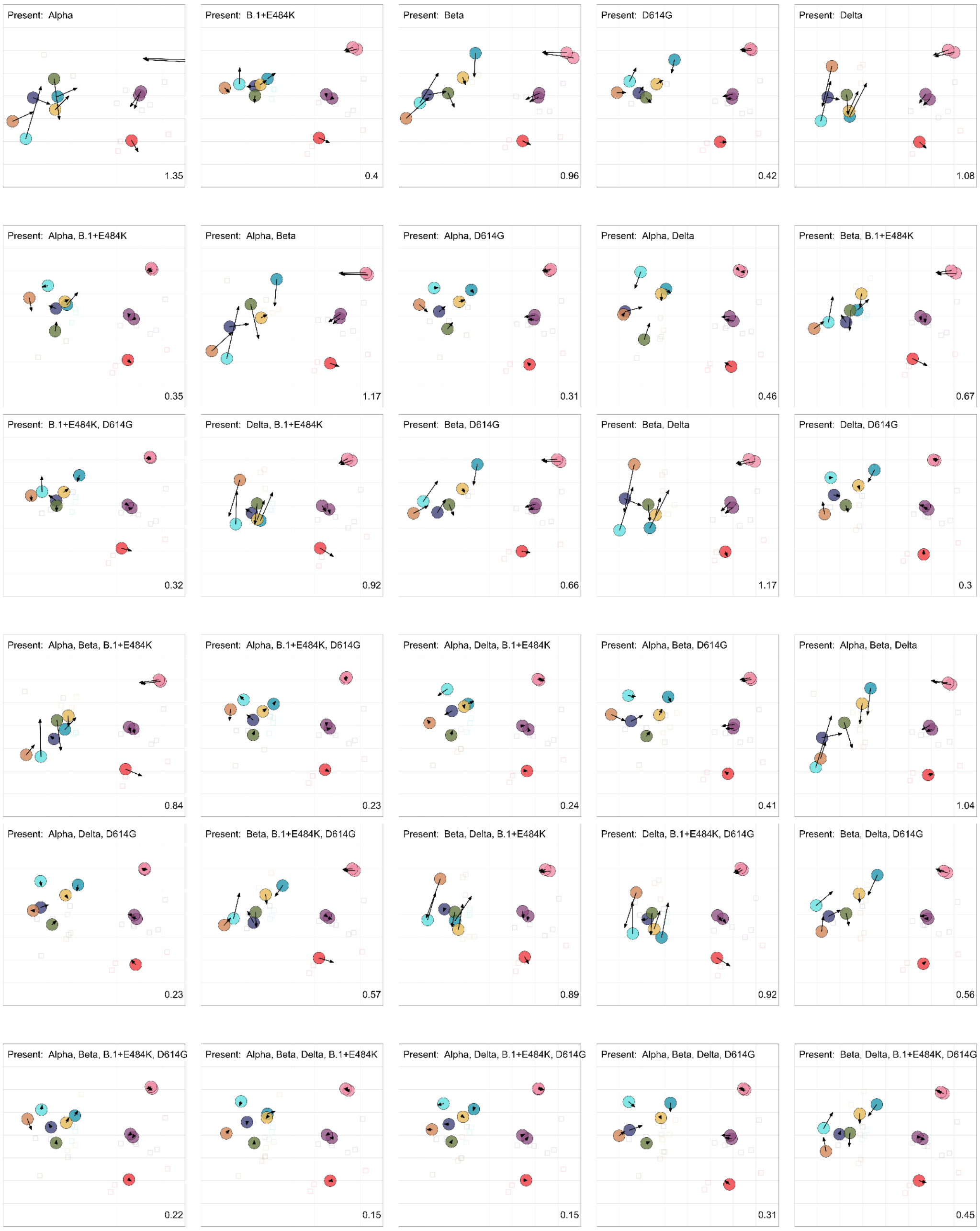

**Figure S26: Subsampling serum groups.** Each map was made from the BA.1 and BA.2 sera, and one (first row), two (second and third row), three (fourth and fifth row), or four (bottom row) pre-Omicron serum groups. The arrows point to the position of the variants in a map made from the all-against-all titrated sera and variants (without the Omicron BF.7, BQ.1.18, XBB.2, and BN.1.3.1 variants, and the BA.5 convalescent sera). The number in the bottom right corner shows the total root mean squared deviation of the positions of the variants between the full map and the subsampled map. The text in the top-left indicates the included serum groups, in addition to the BA.1 and BA.2 serum groups. Each map was made with 100 optimisations.

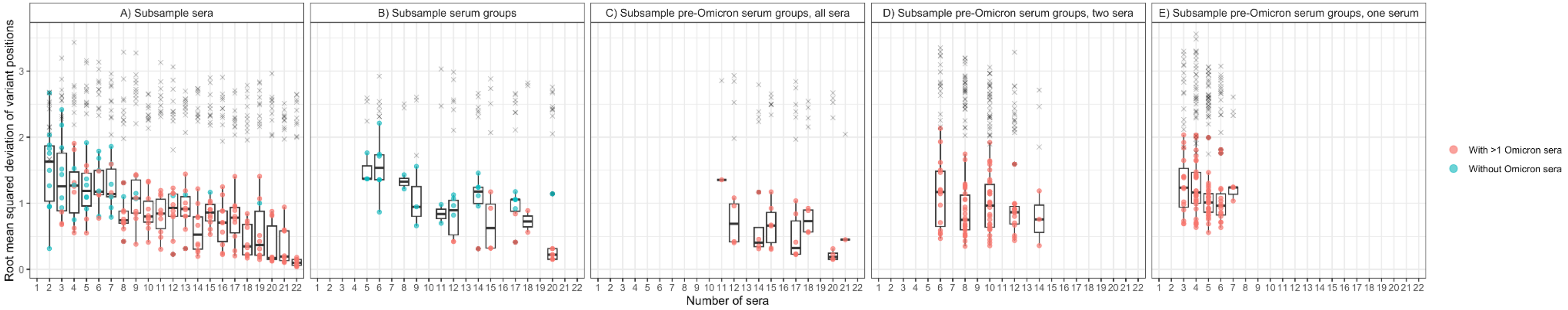

**Figure S27: Root mean squared deviation (RMSD) of variant positions of subsampled maps compared to complete map.** In order to investigate map stability when reducing the number of sera used, we subsampled a map made from the all-against-all titrated sera and variants (without the Omicron BF.7, BQ.1.18, XBB.2, and BN.1.3.1 variants, and the BA.5 convalescent sera) in different ways: A) Randomly subsample from all sera. B) Randomly sample which serum groups are included. C) Randomly sample which serum groups are included, but always include the BA.1 and BA.2 serum groups. D) Randomly subsample the data in panel C to include two sera per serum group. E) Randomly subsample the data in panel C to include one serum per serum group. A map was constructed from each subsample and the RMSD of the variant positions in the subsampled map compared to a map made from the all-against-all titrated sera and variants was calculated. The RMSD is indicated by boxplots and coloured dots, according to the number of sera in each subsampled map. Red dots indicate maps that include both the BA.1 and BA.2 serum groups, blue dots indicate maps that include either the BA.1 or BA.2 or neither of the BA.1 or BA.2 serum groups. To estimate the RMSD under a random scenario, titers in each subsampled map were randomized, a new map constructed and the RMSD from the randomized, subsampled map compared to the full map in Figure 3 was calculated, and is indicated by ‘x’. Subsampled maps were constructed in two dimensions, using 100 optimisations. As the Alpha serum group only includes two sera, and the BA.2 serum groups contain six sera in total, the number of sera in panels B and C is not always a multiple of three.

### Supplementary tables

| Variant | Spike substitutions relative to Wuhan-Hu-1 (EPI_ISL_402125) | Stock-ID | Additional substitutions and minor variants >5% of reads in S/>20% of reads in rest of the genome | GISAID accession number |
|---|---|---|---|---|
| D614G | D614G | SARS-CoV-2 984 p2 vs 12.10.2020; FW (V66) | Synonymous minor variant at nt position 11758 (C>T, 5% of reads), 24616 (C>T, 8% of reads), Non-synonymous minor variant at nt position 23525 (S:H655Y, 6% of reads), 23606 (S:R682W, 6% of reads). | EPI_ISL_406862 (original sample sequence), EPI_ISL_16737908 (PRNT stock sequence) |
| Alpha | H69-, V70-, Y144-, N501Y, A570D, D614G, P681H, T716I, S982A, D1118H | SARS-CoV-2/21528/BaWü p.2 VS 26.03.2021 3dpi, V86 | Non-synonymous substitution at nt position 25511 (ORF3a:S40L), 27533 (ORF7a:H47P), 27731 (ORF7a:C113Y) Synonymous substitution at nt position 6379. | EPI_ISL_754174 (original sample sequence), EPI_ISL_16738010 (PRNT stock sequence) |
| Beta | L18F, D80A, D215G, L241-A243del, K417N, E484K, N501Y, D614G, A701V | SARS-CoV-2/22131/B.1.351 Südafrika p.2, VS 05.02.2021, 3 dpi; AR, V78 | Non-synonymous minor variant at nt position 10533 (3CL:C160F, 9% of reads), 10809 (3CL:P252L 19% of reads), 11454 (ORF1a:A3730S, 5% of reads), 11750 (ORF1a:L3829Y, 24% of reads). Non-synonymous substitution at nt position 23607 (S:R682Q), 25563 (ORF3a:Q57H), 25784 (ORF3a:W131L), 27409 (ORF7a:F6I), Synonymous substitution at nt position 6349, Synonymous minor variant at nt position 5392 (28% of reads) | EPI_ISL_862149 (original sample sequence), EPI_ISL_16738012 (PRNT stock sequence) |
| Delta | T19R, G142D, E156G, F157-, R158-, L452R, T478K, D614G, P681R, D950N | V7_25893_23_C.V.p1 | Non-synonymous minor variant at nt position 23606 (S:R682W, 19% of reads).* | EPI_ISL_3710429 (original sample sequence of V7_25893_23_C.V.p1), EPI_ISL_16465138 (PRNT stock sequence) |
| Omicron BA.1 | L5F, A67V, H69-, V70-, T95I, G142D, V143-Y145del, N211I, L212-, -215E, -216P, -217E, G339D, S371L, S373P, S375F, K417N, N440K, G446S, S477N, T478K, E484A, Q493R, G496S, Q498R, | V1_26335_C.V.p2 | Non-synonymous change in ORF1ab L3829F. Synonymous minor variants at nt positions 17590 and 24382. | EPI_ISL_7019047 (original sample sequence) |

| | | | | |
|---|---|---|---|---|
| | N501Y, Y505H, T547K, D614G, H655Y, N679K, P681H, N764K, D796Y, N856K, Q954H, N969K, L981F | | | EPI_ISL_16359096 (PRNT stock sequence) |
| Omicron BA.2 | T19I, L24S, P25-A27del, G142D, V213G, G339D, S371F, S373P, S375F, T376A, D405N, R408S, K417N, N440K, S477N, T478K, E484A, Q493R, Q498R, N501Y, Y505H, D614G, H655Y, N679K, P681H, N764K, D796Y, Q954H, N969K | V2_26729_12_V.V.p6 | Non-synonymous change in C3L protease P96H. Synonymous minor variant at nt position 11750. Non-synonymous minor variants at 13131 (ORF1ab:Q4289R), 23606 (S:R682W, 16% of reads), 23757 (S:T732I, 7% of reads), 25618 (ORF3a:G76S, 40% of reads), 26461 (E:L73F, 24% of reads) | EPI_ISL_9553935 (original sample sequence) EPI_ISL_16359856 (PRNT stock sequence) |
| Omicron BA.2 | T19I, L24S, P25-A27del, G142D, V213G, G339D, S371F, S373P, S375F, T376A, D405N, R408S, K417N, N440K, S477N, T478K, E484A, Q493R, Q498R, N501Y, Y505H, D614G, H655Y, N679K, P681H, N764K, D796Y, Q954H, N969K | V3_26729_2_V.V.p6 | Non-synonymous change in C3L protease P96H, in E L73F. Non-synonymous minor variant at nt position 2106 (ORF1ab:T614I, 23% of reads), 11355 (ORF1a:A3697V, 23% of reads), 11454 (ORF1a:A3730V 34% of reads), 12789 (ORF1a:T4175I, 29% of reads), 23606 (S:R682W, 8% of reads), 23607 (S:R682L, 16% of reads), 23757 (S:T732I, 34% of reads), 26542 (M:T7I, 28% of reads). | EPI_ISL_9553926 (original sample sequence) EPI_ISL_16359857 (PRNT stock sequence) |
| rSARS-CoV-2_B.1+E484K | E484K, D614G | B.1 + E484K (p1,VS 07.10.2021) from Simon, V217 | Synonymous substitution at nt position 18822. Non-synonymous minor variant at nt position 7493 (ORF1a:M2410L, 26% of reads), 11403 (ORF1a:N3713S), 11522 (ORF1a:F3753V, 7% of reads), 23607 (S:R682P, 19% of reads), synonymous minor variant at nt position 11332 (7% of reads), 29662 (16% of reads) | EPI_ISL_16738016 (PRNT stock sequence) |
| Omicron BA.4 | V3G, T19I, L24S, P25-A27del, H69-, V70-, G142D, V213G, G339D, S371F, S373P, S375F, T376A, D405N, R408S, K417N, N440K, L452R, S477N, T478K, E484A, F486V, Q498R, N501Y, Y505H, D614G, H655Y, N679K, P681H, N764K, D796Y, Q954H, N969K | V38_29061_V.V.p4 | Non-synonymous change in nsp6 A161V | EPI_ISL_16221624 (PRNT stock sequence) |
| Omicron BA.5 | T19I, L24S, P25-A27del, H69-, V70-, G142D, V213G, G339D, S371F, S373P, S375F, T376A, D405N, R408S, K417N, N440K, L452R, S477N, T478K, E484A, F486V, Q498R, N501Y, Y505H, D614G, H655Y, N679K, P681H, N764K, D796Y, Q954H, N969K | V34_29057_V.V.p4 | Non-synonymous change in membrane glycoprotein V221L | EPI_ISL_16221625 (PRNT stock sequence) |

| | | | | |
|---|---|---|---|---|
| Mu | T95I, Y144T, Y145S, -146N, R346K, E484K, N501Y, D614G, P681H, D950N | V44_29349_V.V.p2 | Non-synonymous minor variant at nt positions 23583-23597 (18% of reads with a deletion from aa position S:Y674- to N679-), nt positions 26284-26286 (43% of reads with a deletion of aa position E:V14). Non-synonymous change in ORF3a Q57H, L106F. | EPI_ISL_17566125 (PRNT stock sequence) |
| Omicron BF.7 | T19I, L24S, P25-A27del, H69-, V70-, G142D, V213G, G339D, S371F, S373P, S375F, T376A, D405N, R408S, K417N, N440K, L452R, S477N, T478K, E484A, F486V, Q498R, N501Y, Y505H, D614G, H655Y, N679K, P681H, N764K, D796Y, Q954H, N969K | V66_29620_C.V.p1 | Non-synonymous minor variant (28289, N:P6T, 7% of reads), synonymous substitution (28330, A>G), non-synonymous substitution (29253, N:S327L) | EPI_ISL_17293602 (PRNT stock sequence) |
| Omicron BQ.1.18 | T19I, L24S, P25-A27del, H69-, V70-, G142D, V213G, G339D, R346T, S371F, S373P, S375F, T376A, D405N, R408S, K417N, N440K, K444T, L452R, N460K, S477N, T478K, E484A, F486V, Q498R, N501Y, Y505H, D614G, H655Y, N679K, P681H, N764K, D796Y, Q954H, N969K | V60_40617_u.V.p3 | Synonymous substitutions in ORF1ab (10156 and 18060), ORF3a (25584), synonymous minor variants (S:24814, T>C (55%)), non-synonymous minor variant in ORF6 (D>L, 51% of reads), non-synonymous substitution in E (V14-, 74% of reads with deletion), N (G>-, 86% of reads), N (A50S) | EPI_ISL_17293256 (PRNT stock sequence) |
| Omicron BN.1.3.1 | T19I, L24S, P25-A27del, H69Y, G142D, K147E, W152R, F157L, I210V, V213G, G257S, G339H, R346T, K356T, S371F, S373P, S375F, T376A, D405N, R408S, K417N, N440K, G446S, N460K, S477N, T478K, E484A, F490S, Q498R, N501Y, Y505H, D614G, H655Y, N679K, P681H, N764K, D796Y, Q954H, N969K | V78_41249_V.V.p4 | Synonymous minor variant (4456, ORF1a, C>T, 10% of reads), (28420, N, T>C, 94% of reads), (28558, N, T>A, 67% of reads), Non-synonymous minor variant (10809, C3L proteinase, P252L, 89% of reads), (11750, ORF1a, L3829Y, 8% of reads), (13965, ORF1ab, T4567C, 7% of reads), (14724, ORF1ab, S4820F, 10% of reads), (25337, S, D1259H, 6% of reads), (26261, E, S6L, 6% of reads), (26309, E, A22D, 63% o freads), Non-synonymous substitutions (10809, C3L proteinase, P252L, 89% of reads), (26309, E, A22D, 63% o freads), synonymous substitution (28420, N, T>C, 94% of reads), (28558, N, T>A, 67% of reads) | EPI_ISL_17548230 (PRNT stock sequence) |
| Omicron XBB.2 | T19I, L24S, P25-A27del, V83A, G142D, Y144-, H146Q, Q183E, V213E, D253G, G339H, R346T, L368I, S371F, S373P, S375F, T376A, D405N, R408S, K417N, N440K, V445P, G446S, N460K, S477N, T478K, E484A, F486S, F490S, Q498R, | V79_41245_V.V.p4 | Non-synonymous minor variant (9852, ORF1a, D3196A, 39% of reads), (11634, ORF1a, C3790L, 33% of reads), (11635, ORF1a, C3790E, 37% of reads), (12357, ORF1a, T4031None, 6% of reads), (26369, E, Y42C, 12% of reads), (26284-26286, E, deletion, 76% of reads), Synonymous minor variant (27167, M, C>T, 14% of reads) Non-synonymous substitution (26284-26286, E, deletion, 76% of reads) | EPI_ISL_17549547 (PRNT stock sequence) |

| | |
|---|---|
| | N501Y, Y505H, D614G, H655Y, N679K, P681H, N764K, D796Y, Q954H, N969K |

**Table S1: Description of isolates.** *Re-sequencing of the stock used for hamster infection showed a non-synonymous substitution in S (S:L179F) for this isolate.

| Serum group | Serum | Variant | Original titer | Adapted titer |
|---|---|---|---|---|
| Alpha convalescent | 2.1 | Alpha | 3010.13 | 3795 |
| Alpha convalescent | 2.2 | BA.2 | 830.37 | 640 |
| Beta convalescent | 3.2 | BA.2-12 | <160 | 183.5 |
| Beta convalescent | 3.3 | Alpha | 1372.73 | 968.83 |
| Beta convalescent | 3.3 | Beta | 2057.81 | 1824.34 |
| Delta convalescent | 4.1 | Alpha | >5120 | 5120 |
| Delta convalescent | 4.2 | Delta | 3125.64 | 3570.23 |
| Delta convalescent | 4.3 | Alpha | 2382.62 | 1622.41 |
| Delta convalescent | 4.3 | Delta | >5120 | 5120 |
| BA.1 convalescent | 5.1 | Alpha | 2898.75 | <40 |
| BA.1 convalescent | 5.1 | BA.2-12 | 409.82 | 326.59 |
| BA.1 convalescent | 5.2 | Alpha | 205.25 | 164.89 |
| BA.1 convalescent | 5.2 | BA.2 | 163.89 | 225.11 |
| BA.1 convalescent | 5.3 | BA.2 | <160 | 220.56 |
| BA.2 convalescent-12 | 6.1 | BA.2 | 4699.22 | 5113.91 |
| BA.2 convalescent-12 | 6.2 | BA.2 | >5120 | 5330.82 |
| B.1+E484K convalescent | 7.1 | D614G | >5120 | 5442.78 |
| B.1+E484K convalescent | 7.1 | Alpha | >5120 | 5032.42 |
| B.1+E484K convalescent | 7.1 | Delta | 2116.85 | 2895.08 |

| | | | | |
|---|---|---|---|---|
| B.1+E484K convalescent | 7.1 | B.1+E484K | 2437.81 | 2635.04 |
| B.1+E484K convalescent | 7.1 | BA.2-12 | 2607.2 | 2859.64 |
| B.1+E484K convalescent | 7.1 | Mu | 6364.97 | >5120 |
| B.1+E484K convalescent | 7.2 | Mu | 2419.48 | 2034.42 |
| B.1+E484K convalescent | 7.3 | Alpha | 2962.97 | 4540.72 |
| B.1+E484K convalescent | 7.3 | Mu | 5237.84 | >5120 |
| D614G convalescent | 8.1 | BA.2-12 | 557.5 | 320 |
| BA.5 convalescent | 9.1 | BF.7 | 1922.55 | 1647.68 |
| BA.5 convalescent | 9.3 | BQ.1.18 | 2909.38 | 3539.07 |

**Table S2: Adaptations made to the PRNT90 titers inferred with constraining the neutralization curve at zero.** In the titrations above, visual inspection of the titer curve revealed that the neutralization curve fixed at zero did not give an ideal fit. Therefore, the adaptations listed above were made to those titers.

4. R interface to CmdStan, (available at https://mc-stan.org/cmdstanr).

5. Stan. *stan-dev.github.io*, (available at https://mc-stan.org).
